# DNMT3A-dependent DNA methylation is required for spermatogonial stem cells to commit to spermatogenesis

**DOI:** 10.1101/2021.04.19.440465

**Authors:** Mathilde Dura, Aurélie Teissandier, Mélanie Armand, Joan Barau, Lorraine Bonneville, Michael Weber, Laura G. Baudrin, Sonia Lameiras, Deborah Bourc’his

## Abstract

DNA methylation plays a critical role in spermatogenesis, as evidenced by the male sterility of DNA methyltransferase (DNMT) mutant mice. Here, we report a striking division of labor in the establishment of the methylation landscape of male germ cells and its functions in spermatogenesis: while DNMT3C is essential for preventing retrotransposons from interfering with meiosis, DNMT3A broadly methylates the genome—at the exception of DNMT3C-dependent retrotransposons—and controls spermatogonial stem cell (SSC) plasticity. By reconstructing developmental trajectories through single-cell RNA-seq and by profiling chromatin states, we found that *Dnmt3A* mutant SSCs can only self-renew and no longer differentiate due to spurious enhancer activation that enforces an irreversible stem cell gene program. We therefore provide a novel function for DNA methylation in male fertility: the epigenetic programming of SSC commitment to differentiation and to life-long spermatogenesis supply.

## Introduction

Cytosine DNA methylation is an epigenetic mark that is crucial for proper mammalian development. Promoter methylation provides stable and long-term repression, with little variation in patterns across somatic tissues. This regulatory mode mostly applies to retrotransposon control, and to the constitutive repression of a small subset of genes, such as germline genes, and genes subject to genomic imprinting or X chromosome inactivation^1, 2^. By contrast, enhancer methylation is prevalent and dynamic, across tissues and developmental stages^3–5^. The identification of methyl-sensitive transcription factors (TFs) in biochemical assays has provided a conceptual frame for the function of enhancer DNA methylation^6, 7^. However, there is still limited *in vivo* evidence that DNA methylation modulates gene programs by limiting TF-enhancer functional interactions during differentiation processes.

Male germline differentiation provides a highly relevant context to study the breadth of DNA methylation distribution and function. Germline epigenetic reprogramming produces an extensively hypomethylated genome, onto which male germ cell-specific DNA methylation is established during fetal life, and impacts all genomic compartments: genes, intergenic sequences and retrotransposons. Remethylation occurs prior to the formation of spermatogonial stem cells (SSCs), which sustain life-long spermatogenesis through their dual capacity to self-renew and differentiate^8^. Incidentally, DNA methylation patterns established in fetal germ cells are mostly unaltered in post-natal life, and are propagated from SSCs to the successive differentiating types that lead to spermatozoa production^9^.

Male germline methylation requires two *de novo* methyltransferases in mice, DNMT3A and DNMT3C, and a catalytically inactive co-factor, DNMT3L^1^. Individual mutations in these genes consistently lead to male sterility, highlighting the key role of DNA methylation for spermatogenesis^10–12^. The recently discovered DNMT3C enzyme selectively methylates the promoters of young retrotransposon lineages, which represents only 1% of the mouse genome^12^. Nevertheless, this restricted function is absolutely essential for meiosis: failure to establish retrotransposon methylation in fetal stages results in their post-natal activation at meiosis, subsequent perturbation of the meiotic chromatin landscape and spermatogenic interruption by apoptosis, as observed in *Dnmt3C* and *Dnmt3L* mutants^12–14^. Prior to the identification of DNMT3C, DNMT3A was regarded as the sole enzyme responsible for male germline methylation, including at retrotransposons. This idea is still persistent although DNMT3A targets have never been profiled genome-wide during germline reprogramming and the nature of the spermatogenetic impairment of *Dnmt3A* mutants remains uncertain^11, 15, 16^.

Here, we demonstrate that DNMT3A function in male germline development is not related to retrotransposon control and meiosis protection. Rather, DNMT3A-dependent DNA methylation pre-emptively programs the capacity of spermatogonial stem cells to commit to spermatogenetic differentiation after birth, by limiting aberrant enhancer activity associated with stem cell identity.

## Results

### DNMT3A broadly methylates the genome in fetal male germ cells

DNMT3A was previously shown to methylate paternally imprinted genes in male germ cells^11, 16^. However, contrary to the DNMT3L co-factor and the retrotransposon-specific DNMT3C enzyme, there is currently no genome-wide maps of DNMT3A targets during male germ cell development. We therefore performed Whole-Genome Bisulfite Sequencing (WGBS) on DNA extracted from wildtype (WT) and *Dnmt3A* mutant (*Dnmt3A* knockout, *Dnmt3A^KO^*) prospermatogonia at embryonic day 18.5 (E18.5) (**Fig. 1a** and **Supplementary Table 1**). Prospermatogonia were isolated by FACS using the *Oct4-eGFP* transgenic line^17^ and libraries were generated by post-bisulfite adaptor tagging (PBAT)^18^. We included age-matched *Dnmt3L^KO^* and *Dnmt3C^KO^* prospermatogonia to allow for direct comparison, as previous whole-genome methylation profiles for these mutants were generated in postnatal germ cells^12^. We chose the E18.5 time-point because *Dnmt3A, 3L* and *3C* genes are upregulated^12, 19^ and *de novo* DNA methylation of male germ cells is still ongoing yet close to completion (**Fig. 1a**), avoiding potential compensatory or secondary effects that could occur later, in postnatal life.

**Fig. 1.**
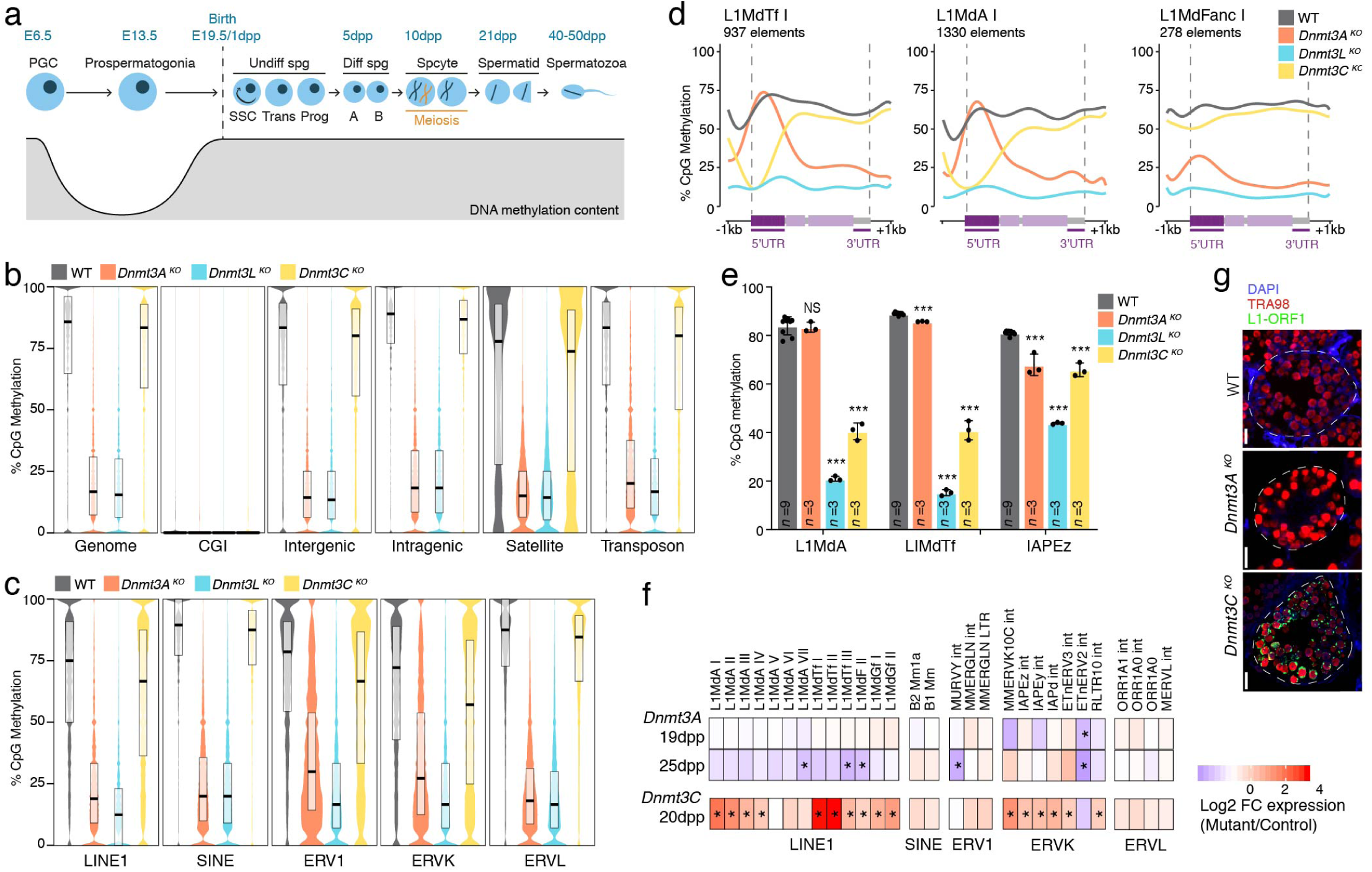
DNMT3A broadly methylates the male germ cell genome but is dispensable for retrotransposon silencing. **a,** Schematic representation of developmental and DNA methylation dynamics of the male germline in the mouse. Postnatal ages indicate predominant germ cell types present in the testis during the first wave of spermatogenesis. PGC: primordial germ cell, SSC: spermatogonial stem cell,Spg: spermatogonia,Trans: transitory, Prog: progenitors. **b,** c,Violin plot representation of CpG methylation content over the whole genome and different genomic compartments (b) and over retrotransposon families (c)in WT (grey), *Dnmt3A^KO^* (orange), *Dnmt3L^K0^* (blue) and *Dnmt3C^K0^* (yellow) fetal germ cells (E18.5), as determined by WGBS. Black horizontal bars represent the median, upper and lower hinges correspond to 75 and 25% quantile. **d,** Metaplots of DNA methylation levels over uniquely assigned full-length elements (>5kb) from three L1 families. e, CpG methylation levels assessed by bisulfite-pyrosequencing of the promoters of indicated retrotransposon families in 1Odpp sorted germ cells. Data are mean ± SD (black bar) from biological replicates, black dots represent biological replicates,n= number of animals (student t-test over WT,*p<0.05,**p<0.005, ***p<0.0005). **f,** RNA-seq heatmap shows log2-fold change (FC) in retrotransposon mRNA levels in *Dnmt3A^KO^* versus WT littermates at 19 and 25dpp (top and middle rows), and previously published changes in *Dnmt3C* mutants versus heterozygous littermates at 20dpp (bottom row)^12^.Annotations are from RepeatMasker. Two biological replicates were sequenced per age and genotype.**g,** L1-0RF1 immunostaining on testis sections at 19dpp in WT,*Dnmt3A^KO^* and *Dnmt3C^K0^* germ cells (TRA98-positive). White dotted lines:tubule delineation. Scale, 20 µm.

Global CpG methylation levels were dramatically reduced in *Dnmt3A^KO^* compared to WT prospermatogonia, with mean values dropping from 67.7% to 20.2%, in an extent similar to *Dnmt3L^KO^* (18.7%) (**Fig. 1b**). We identified 555,893 differentially methylated regions (DMRs), all reflecting hypomethylation in *Dnmt3A^KO^*. By contrast, levels were barely diminished in *Dnmt3C^KO^* (64.5%)—matching previous reports in postnatal germ cells^12^—with a number of 7,620 DMRs (all hypomethylated compared to WT) that did not overlap with *Dnmt3A^KO^* DMRs. DNMT3A appeared necessary for methylating all genomic compartments, including genes, intergenic sequences, and transposable elements, akin to DNMT3L (**Fig. 1b** and **Extended data Fig. 1a**). Both LTR (ERV1, ERVK and ERVL) and non-LTR (LINE1 and SINE) retrotransposons were globally hypomethylated in *Dnmt3A^KO^*, with a reduction 2- to 3- fold greater than observed in *Dnmt3C^KO^*, even at ERVK and LINE1 which are preferential targets of DNMT3C^12^ (**Fig. 1c**). However, metaplot analysis of uniquely mapped copies from individual retrotransposon families revealed a striking inverted pattern of defective DNA methylation in *Dnmt3A^KO^* and *Dnmt3C^KO^* germ cells. When considering evolutionarily young L1 families (L1Md-TfI and L1Md-AI), methylation of the promoters of these elements was confirmed to be DNMT3C-dependent uniquely, while DNMT3A was required for methylating the entire body of these same elements (**Fig. 1d** and **Extended data Fig. 1a**). In evolutionarily old L1s (L1Md-FancI), DNMT3A was responsible for methylating the full element length, while DNMT3C was generally dispensable. This situation was similar for ERVL elements (**Extended data Fig. 1b**), while uniquely assigned ERVK elements belonging to the young IAPEz family relied on DNMT3C only (**Extended data Fig. 1c**). In contrast with these specificities, DNMT3L had a global role in assisting *de novo* DNA methylation in fetal male germ cells: *Dnmt3L^KO^* prospermatogonia displayed hypomethylation over both DNMT3A and DNMT3C genomic targets (**Fig. 1d** and **Extended Fig. 1a-c**).

Overall, we reveal here that DNMT3A is a largely indiscriminate enzyme that *de novo* methylates the whole genome of prenatal male germ cells, with the notable exception of young retrotransposon promoters (**Extended data Fig. 1d**).

### DNMT3A does not silence retrotransposons during spermatogenesis

*Dnmt3A^KO^* fetal germ cells display normal methylation at young retrotransposon promoters, which implies proper silencing. However, *Dnmt3A* mutants were previously reported to phenocopy *Dnmt3L* mutants^11^, in which retrotransposon reactivation culminates after birth in meiotic cells, in association with apoptosis and spermatogenesis interruption^13, 14^. We therefore went on to verify whether defective retrotransposon methylation and silencing could occur postnatally in *Dnmt3A^KO^* germ cells, around meiosis (**Fig. 1a**). Using targeted bisulfite pyrosequencing, we measured promoter DNA methylation of young retrotransposons (L1MdA, L1MdTf and IAPEz) in FACS-sorted germ cells (EpCAM positive; β2-Microglobulin negative) from males at 10 days post-partum (10dpp). At young L1 promoters—similarly to fetal stages—*Dnmt3A^KO^* displayed normal methylation, while *Dnmt3C^KO^* and *Dnmt3L^KO^* showed decreased CpG methylation (**Fig. 1e**). We observed a slight decrease on IAPEz promoters in *Dnmt3A^KO^* (67.7% versus 81.1% in WT), indicating that DNA methylation of a subset of IAP copies may require DNMT3A.

However, no IAP or L1 reactivation was detected in testes of *Dnmt3A^KO^* males (between 19 and 25dpp), at the RNA level using RNA-seq (**Fig. 1f**) and RT-qPCR (**Extended data Fig. 1e**), and at the protein level by immunodetection of L1-encoded ORF1 proteins (**Fig. 1g**), in striking contrast to *Dnmt3C^KO^* and *Dnmt3L^KO^* males. Moreover, SINE and ERVL, which are exclusively methylated by DNMT3A (**Fig. 1c**), also maintained repression in *Dnmt3A^KO^* testes (**Fig. 1f** and **Extended data Fig. 1e**). Lack of retrotransposon reactivation did not reflect a lack of meiotic cells: meiotic genes showed similar mRNA levels in *Dnmt3A* mutants compared to *Dnmt3C* mutants (**Extended data Fig. 1f**). As a whole, these findings definitely exclude a role for DNMT3A in silencing retrotransposons during spermatogenesis. They also allude to the possibility that although they all share a sterility phenotype, the etiology of this sterility may be different in *Dnmt3A* mutants compared to *Dnmt3C* and *Dnmt3L* mutants.

### *Dnmt3A* mutants only progress through the first wave of spermatogenesis

To uncover the function of DNMT3A-dependent DNA methylation in spermatogenesis, we phenotyped *Dnmt3A* mutant testes across ages. Constitutive *Dnmt3A^KO^* animals are developmentally delayed after birth and die around 25dpp^20^ (**Extended data Fig. 2a**). All analyses past 25dpp were therefore performed on germ-cell conditional *Dnmt3A^KO^* by crossing the *Dnmt3A2lox* line with the *Prdm1-Cre* line, which promotes recombination at E9.5^21^ (*Prdm1-Dnmt3A^cKO^*) (**Extended data Fig. 2b**).

From 10dpp to 6 months, *Dnmt3A* mutant males showed significant and increasing reduction in testis weight and seminiferous tubule surface in comparison to their WT littermates (**Extended data Fig. 2c,d**). We then performed histological assessment of testis sections at 19dpp, 6 weeks, 8-9 weeks and 6 months, and compared to age-matched *Dnmt3L^KO^* and *Dnmt3C^KO^* males (**Fig. 2a** and **Extended data Fig. 2e**). As previously reported^12, 13^, *Dnmt3L^KO^* and *Dnmt3C^KO^* tubules exhibited pyknotic nuclei from apoptotic meiotic cells, and lack of subsequent post-meiotic stages. In contrast, we did not detect apoptotic cells in *Dnmt3A* mutant testes, indicating that spermatogenesis might not be interrupted at meiosis. Then, while *Dnmt3C^KO^* tubules continuously produced pre-meiotic cells, *Dnmt3L^KO^* showed progressive spermatogenic decline with age, with only spermatogenesis-free tubules remaining at 6 months. Overall, *Dnmt3A^KO^* and *Prdm1-Dnmt3A^cKO^* mutants displayed a *Dnmt3L^KO^*-like phenotype of progressive loss of spermatogenic ability (**Fig. 2a**).

**Fig. 2.**
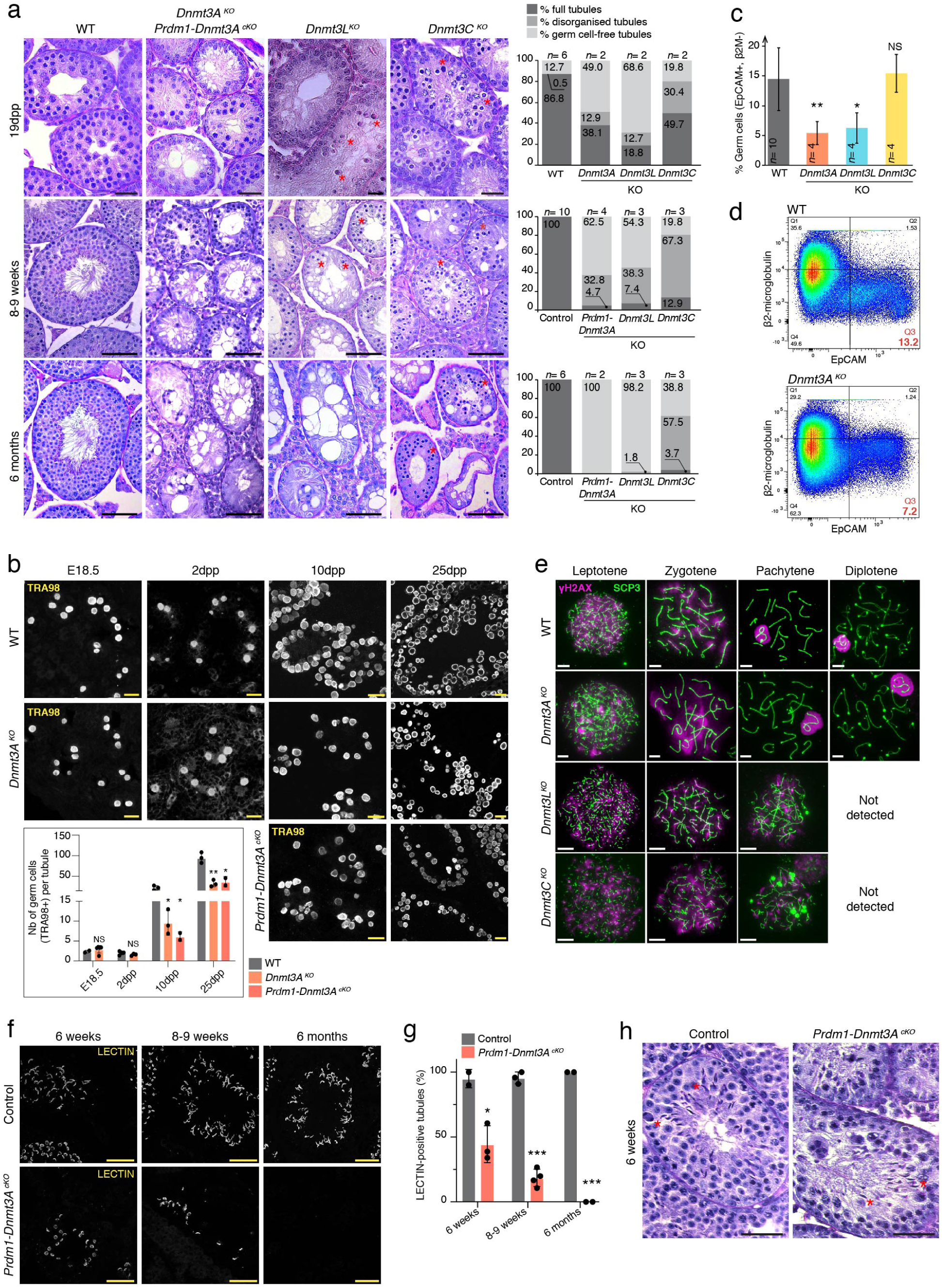
Dnmt3A mutants males complete the first wave of spermatogenesis with reduced germ cell numbers. **a,** (Left) Representative images of Periodic Acid Shift (PAS)-stained testis sections of different genotypes at 19dpp (scale,20 µm), 8-9 weeks and 6 months (scale, 50 µm). Red stars (*) point apoptotic cells. (Right) Quantification of the percentage of different classes of tubules per genotype.n= number of animals. Control genotypes at 8-9 weeks were *Dnmt3A^2lox/KO^; Prdm1-Cre^0/0^* (n= 2), *Dnmt3A^KO/WT^; Prdm1-Cre^Tg/O^, Dnmt3A^2lox/WT^; Prdm1-Cre^0/0^, Dnmt3L^KO/WT^* (n= 3) and *Dnmt3C^KO/WT^* (n= 3). Control genotype at 6 months were *Dnmt3A^cKO^* (n= 4), *Dnmt3A^21ox/WT^; Prdm1-Cre^0/0^* and *Dnmt3A^KO/WT^; Prdm1-Cre^Tg/O^.* **b,** (Top) Representative image of TRA98 staining on testis sections from WT, *Dnmt3A^KO^* and *Prdm1-Dnmt3N^K0^* males at E18.5,2dpp, 10dpp and 25dpp (scale,20 µm).(Bottom) Quantification ofTRA98-posi­ tive cells normalized per tubule.Data are mean ± SD (black bar) and individual points represent biological replicates (student t-test over WT *p<0.05, **p<0.005). c, Percentage of live germ cells (DAPl-Neg, EpCAM-Pos, f32M-Neg) per testis of different genotypes at 1Odpp,assessed by FACS. Data are mean ± SD (black bar) from at least four biological replicates,n= number of animals (student t-test over WT, NS= nonsignificant, *p<0.05, **p<0.005). **d,** Representative FACS plots of live EpCAM-Pos, f32M-Neg gated testicular cells. from 10dpp *Dnmt3A^KO^* and WT males. Cell percentages are indicated in each quarter,with percentage of EpCAM-Pos, 2M-Neg cells in red. e, Representative microscopy images of SCP3 (green) and yH2AX (pink) immunodetection on WT, *Dnmt3A^KO^, Dnmt3L^K0^* and *Dnmt3C^K0^* meiotic spreads (16-25dpp). Diplotene stages were absent in *Dnmt3L^K0^* and *Dnmt3C^K0^* Scale, 5 µm. f, Representative image of LECTIN (acrosome/haploid marker) staining on testis sections from *Prdm1-Dnmt3N^K0^* and controls at indicated ages. Scale, 50 µm. g, Quantification of tubule percentage with LECTIN-stained cells. Control genotypes at 6 weeks : *Dnmt3A^KO/WT^; Prdm1-Cre^Tg/0^* and *Dnmt3A^2lox/KO^; Prmd1-Cre^0/0^* - at 8-9 weeks : *Dnmt3A^2lox/KO^; Prmd1-Cre^0/0^* (n= 2) and *Dnmt3A^2lox/WT^; Prdm1-Cre^0/0^* - at 6 months: *Dnmt3A^KO/WT^; Prmd1-Cre^Tg/0^* and *Dnmt3A^2lox/WT^; Prmd1-Cre^0/0^.* Data are mean ± SD (black bar) and individual points represent biological replicates (student t-test over WT *p<0.05, **p<0.005, ***p<0.0005). h, Representative images of PAS-stained testis sections of 6 week-old *Prdm1-Dnmt3N^K0^* male and control littermate *(Dnmt3A^2^1ox1Ko; Prdm1-Cre^0/0^).* Red stars (*) point spermatozoa heads. Scale, 30 µm.

To pinpoint the onset of spermatogenic failure, *Dnmt3A* mutant germ cells were counted across the first wave of spermatogenesis (around birth time to 6 weeks) (**Fig. 1a**), by immunodetection of the pan germ cell marker TRA98 (**Fig. 2b**). No difference was scored compared to WT immediately before (E18.5) and after birth (2dpp): *Dnmt3A^KO^* males were therefore born with appropriate numbers of prospermatogonia, from which the postnatal spermatogonial contingent emerges. However, starting at 10dpp, germ cell depletion became significant, with a 2- to 3-fold reduction in average numbers per tubule compared to age-matched WT. This was confirmed by FACS analysis (**Fig. 2c, d**): 5% of testicular cells were scored as “EpCAM-pos; β2M-neg” germ cells in 10dpp-old *Dnmt3A^KO^* males against 15% in WT. This reduction rate was similar in *Dnmt3L^KO^* testes while *Dnmt3C^KO^* had normal germ cell counts. Importantly, germ cell depletion was an intrinsic cell defect, being also observed in germ cell-specific *Prdm1-Dnmt3A^cKO^* mutants (at 10dpp, **Fig. 2b**).

Consistent with the lack of apoptotic cells in histological sections (**Fig. 2a**), prepuberal *Dnmt3A^KO^* males completed the first prophase of meiosis: diplotene figures were readily detectable upon SCP3 and γH2AX labeling of meiotic chromosome spreads around 20dpp (**Fig. 2e** and **Extended data Fig. 2f, g**). This was in stark contrast to *Dnmt3C^KO^* and *Dnmt3L^KO^* males where spermatogenesis was arrested prior to pachytene (**Fig. 2e**). RT-PCR detection of haploid cell markers further indicated that spermatids were specified in constitutive *Dnmt3A^KO^* males at 25dpp (**Extended data Fig. 2h**). Lectin-mediated staining of acrosome structures confirmed the presence of round and elongated spermatids in testes of *Prdm1-Dnmt3A^cKO^* males at 6 weeks—with half numbers of lectin-positive tubules compared to controls—(**Fig. 2f, g**) and morphologically mature spermatozoa were detected (**Fig. 2h** and **Extended data Fig. 2i**). However, as described above (**Fig. 2a**), pre- and post-meiotic germ cells eventually disappeared from *Prdm1-Dnmt3A^cKO^* testes, upon initiation of subsequent rounds of spermatogenesis, and were visibly absent at 6 months (**Fig. 2f,g** and **Extended data Fig. 2f**).

In sum, contrary to *Dnmt3C* and *Dnmt3L* mutants, *Dnmt3A* mutants*—*both constitutive and germ cell-conditional*—*can progress past meiosis. This likely relates to the lack of retrotransposon reactivation in *Dnmt3A* mutants. However, *Dnmt3A* mutants can only fulfill the first postnatal wave of spermatogenesis, after which they lose spermatogenic potential. Interestingly, the first cells that enter spermatogenesis—3 to 5 days after birth— emanate from a pool of fetal prospermatogonia that bypass the spermatogonial stem cell (SSC) state and directly transition to differentiated spermatogonia^22^ (**Fig. 1a**). Meanwhile, the remainder of prospermatogonia generate the foundational SSC compartment, from which spermatogenesis initiates throughout the reproductive lifespan^23^. The phenotypic features of *Dnmt3A* mutants suggests that SSC-dependent spermatogenesis is compromised.

### SSCs accumulate in *Dnmt3A* mutants

We next wanted to determine whether SSC establishment, self-renewal or commitment to differentiation was altered in *Dnmt3A* mutants. By counting GFRA1-positive cells (a general SSC marker) on testis sections, we excluded a problem in the initial establishment of the SSC pool in *Dnmt3A^KO^* neonates: at 10dpp, similar numbers of GFRA1-positive cells were observed compared to WT littermates (**Fig. 3a**). Unexpectedly, at all subsequent ages examined, we did not observe progressive exhaustion of the SSC pool, but rather increased GFRA1-positive cells, in both constitutive and conditional *Dnmt3A* mutants compared to control mice (**Fig. 3a**), while somatic Sertoli cells (SOX9-positive) remained in normal amounts (**Extended data Fig. 3a,b**). In control mice, the number of GFRA1-positive cells per tubule mm^2^ declined by 30-fold from 10dpp to 6 weeks; comparatively, it dropped by 5-fold only in *Dnmt3A* mutants (**Fig. 3a**). In 6 week-old *Prdm1-Dnmt3A^cKO^* males, GFRA1-stained cells represented 3.5% of all germ cells (VASA positive), versus 1.2% in control mice (**Fig. 3b**). At 6 months, although *Prdm1-Dnmt3A^cKO^* tubules appeared as germ cell-free by histological examination (**Fig. 2a**), GFRA1-positive cells were actually still present, at least 3-times more abundantly than in tubules of control mice (**Fig. 3a**), and were the quasi-exclusive germ cell type to remain (85%) (**Fig. 3b**). Comparatively, GFRA1-positive cells are rare in control males at this age, representing 0.9% of all germ cells. To better define the origin of this potential SSC increase, we utilized the *Id4-eGFP* reporter transgenic line^24, 25^, which we crossed onto the *Dnmt3A^KO^* background (**Extended data Fig. 3b**). The *Id4-eGFP* transgene allows sorting undifferentiated spermatogonia sub-populations along a continuum that ranges from “ID4-GFP^bright^” enriched in regenerative SSCs at the top of the SSC hierarchy to “ID4-GFP^dim^” enriched in non-regenerative spermatogonial progenitors; “ID4-GFP^medium^” denotes various intermediate and transitory states^25^ (**Fig.1a**). FACS analysis at 10dpp revealed a 10-fold excess in ID4-GFP^bright^ SSCs in *Dnmt3A^KO^* compared to WT testes (19.45% versus 2.27% of all ID4-GFP-positive cells) and a 3-fold excess in ID4-GFP^medium^, while ID4-GFP^dim^ progenitors were relatively reduced by half (**Extended data Fig. 3c**).

**Fig. 3.**
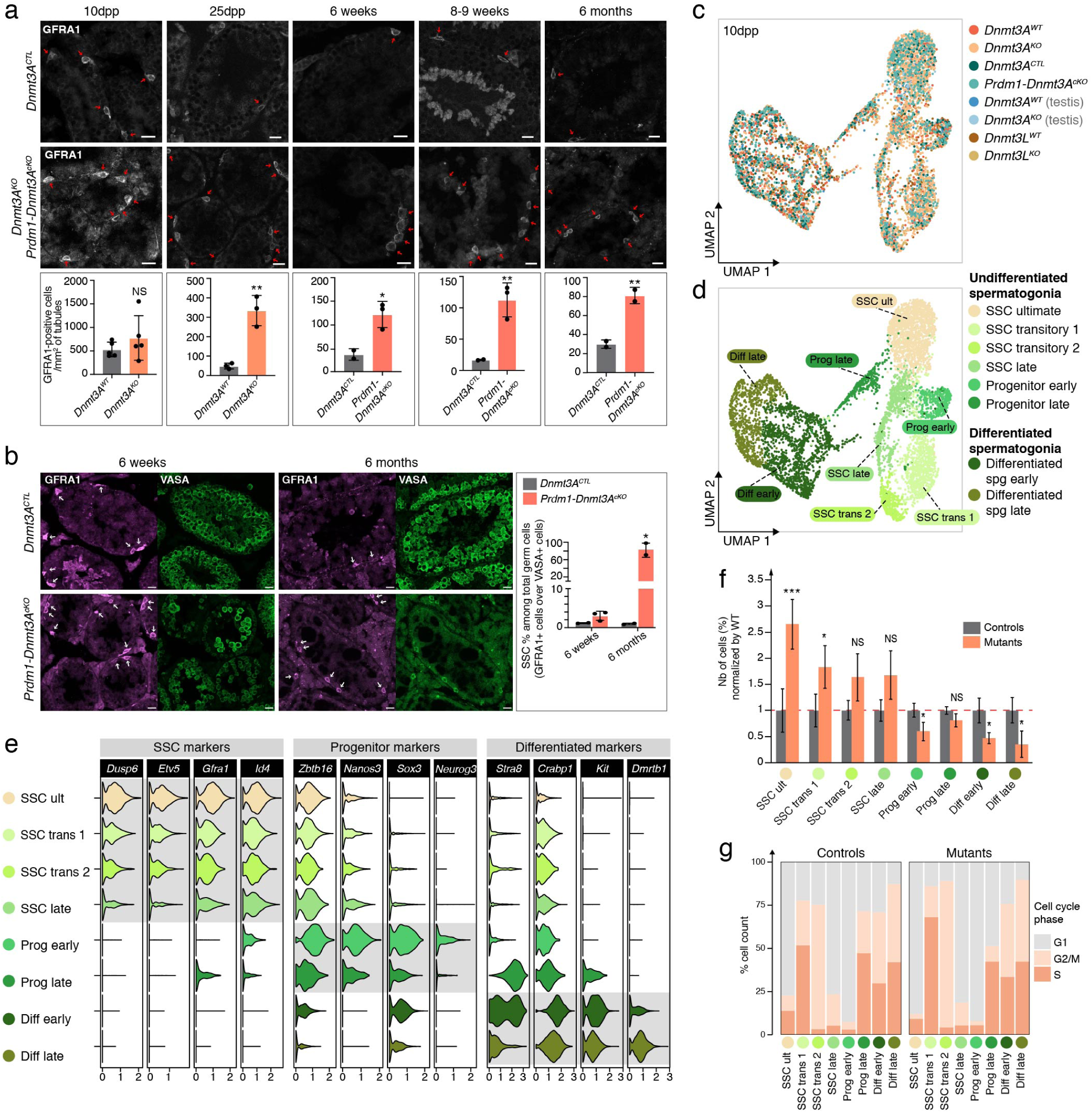
SSCs accumulate in absence of DNMT3A-dependent DNA methylation. **a,** (Top) Representative image of GFRA1 (SSC marker) staining on testis sections from *Dnmt3A^KO^, Prdm1-Dnmt3Ac^K0^* and controls at indicated ages. Scale, 20 µm. (Bottom) Quantification of GFRA1-positive cells per mm2 of tubule. Control genotypes at 6 weeks: *Dnmt3A^KO/WT^; Prdm1-CreTg/O* and *Dnmt3A^2lox/KO^; Prmd1-Cre^0/0^* - at 8-9 weeks: *Dnmt3A^2lox/KO^; Prmd1-Cre^0/0^* (n=2) and *Dnmt3A^2lox/WT^; Prdm1-Cre^0/0^* - at 6 months: *Dnmt3^KO/WT^; Prmd1-Cre^Tg/O^* and *Dnmt3A^2lox/WT^; Prmd1-Cre^0/0^* Data are mean ± SD (black bar) and individual points represent biologicalreplicates (student t-test over WT, NS:non-significant, *p<0.05,**p<0.005). **b,** (Left) Representative microscopy images of GFRA1 (purple) and VASA (green) immunodetection at 6 weeks and 6 months. (Right) Quantification of percentage of GFRA1-positive cells (SSCs) among VASA-positive cells (germ cells).Control genotypes at 6 weeks: *Dnmt3^KO/WT^; Prdm1-Cre^Tg/O^* and *Dnmt3A^2lox/KO^ Prmd1-Cre^0/0^* - at 6 months: *Dnmt3A^KO/WT^; Prmd1-Cre^Tg/O^* and *Dnmt3A^2lox/WT^; Prmd1-Cre^0/0^.* Data are mean ± SD (black bar) and individual points represent biological replicates (student t-test over WT *p<0.05). **c,** UMAP dimensionality reduction representation of scRNA-seq-integrated data from 1Odpp germ cells (n= 6,638).Colors represent 8 genotypes and conditions. All conditions correspond to FAGS-enriched germ cells except for “(testis)”, which indicates that all testicular cells were analyzed. **d,** Unbiased cell clustering onto an UMAP representation of scRNA-seq data demonstrated eight germ cell clusters (from cluster 0 to 13 of unbiased cell clustering in Extended data Fig. 5a). **e,** Violin plots show mRNA level variation of different markers of SSCs, progenitors and differentiated spermatogonia among the eight germ cellclusters. **f,** Bar plot showing changes in germ cell cluster frequencies for pooled mutants normalized by controls. Data are mean ± SD (black bar) from four mutant samples *(Dnmt3A^KO^ Prdm1-Dnmt3A^cKO^ Dnmt3^KO^* FACS-enriched germ cells and *Dnmt3A^KO^* all testicular cells) and four control samples *(Dnmt3A^WT^, Dnmt3A^CTL^, Dnmt3L^WT^* FAGS-enriched germ cells and *Dnmt3A^WT^* all testicular cells). Statistical comparison was performed by student t-test over WT, NS= non-significant, *p<0.05, **p<0.005, ***p<0.005. **g,** Comparison of cell cycle signature for controls and mutants for each germ cell cluster.

Therefore, despite the progressive spermatogenic decline of *Dnmt3A* mutants, we found that their SSCs are specified, and can further self-renew and maintain their most naive state, although excessively. We started considering that commitment to differentiation may be defective in *Dnmt3A* mutant SSCs.

### SSCs cannot exit the stem cell state in absence of DNMT3A-dependent DNA methylation

To resolve the origin of the SSC phenotype, we generated unbiased droplet-based single-cell RNA-seq (10X Genomics Chromium) at 10dpp. Eight samples were sequenced, from FACS-enriched EpCAM-pos germ cells (*Dnmt3A^KO^*, *Prdm1-Dnmt3A^cKO^*, *Dnmt3L^KO^* and respective littermate controls) and from whole testis cell suspension (*Dnmt3A^KO^* and WT) (**Supplementary Table 1**). A total of 41,582 cells were mixed, integrated (to reduce noise and variability between conditions) and analyzed together as biological replicates to gain statistical confidence (**Extended data Fig. 4a**). Based on the expression of known markers, we identified germ cells and testicular somatic cells (**Extended data Fig. 4b,c** and **Supplementary Table 2**) and a stringent filtering pipeline allowed us to retain 6,638 high quality single germ cell transcriptomes for analysis (See Methods) (**Fig. 3c** and **Extended data Figs. 4d-f and 5b-d**). Unbiased cell clustering revealed the existence of eight germ cell populations projected into UMAP representation (**Fig. 3d**), whose identities were derived from published classification^25, 26^, known cell-specific markers^8, 27, 28^ and hierarchical clustering: SSC ultimate (SSC ult), SSC transitory 1 and transitory 2 (SSC trans 1 and SSC trans 2), SSC late (SSC late), Progenitor early, Progenitor late, Differentiated spermatogonia early and late (Diff early and Diff late) (**Fig. 3e** and **Extended data Fig. 5e-g**).

The relative frequency of germ cell clusters was affected in mutant samples, with increased SSC populations while progenitors and differentiated spermatogonia were under-represented (**Fig. 3f**). According to our defined criteria, SSC ultimate—the stem cell category at the foundation of spermatogenesis^8, 25^—was the most strongly expanded in mutants, by 3-fold more abundant than in controls (2.7% versus 1%). Importantly, this cellular phenotype was similarly observed when constitutive *Dnmt3A^KO^* and germ cell-conditional *Prdm1-Dnmt3A^cKO^* were considered individually (**Extended data Fig. 5h**), again confirming that the phenotype is germ-cell autonomous. This phenotype could result from failure to acquire DNMT3A-dependent DNA methylation patterns in fetal prospermatogonia and/or from an intrinsic function of DNMT3A in post-natal SSCs, as *Dnmt3A* is expressed in SSCs and their derivatives (**Extended data Fig. 5i**). However, the latter option was disproved by the fact that *Dnmt3L^KO^* mutants also showed SSC expansion by scRNA-seq (**Extended data Fig. 5h**), while *Dnmt3L* is exclusively expressed in fetal prospermatogonia but ceases expression at birth^19, 29^ (**Extended data Fig. 5h,i**). What *Dnmt3A* and *Dnmt3L* mutants have in common is a lack of DNMT3A-dependent DNA methylation inherited from earlier stages, the fetal prospermatogonia, when DNMT3A and DNMT3L are both expressed and establish male germline DNA methylation.

As a first attempt to explain this abnormal cellular distribution, we performed cell cycle analysis^30^. The categories “SSC ultimate”, “SSC late” and “Progenitors early” were the most enriched in G1 phase, suggesting a quiescent state, while “SSC trans”, “Progenitors late” and subsequent categories were mostly cycling cells, as described recently^31^ (**Fig. 3g**).

However, mutant and control samples showed similar cell cycle distribution, excluding that mutant SSC expansion—and “SSC ultimate” notably—results from higher mitotic rate and proliferation. Focusing on *Dnmt3A* mutants specifically (constitutive and germ-cell specific), we then analyzed cellular trajectories in pseudotime using Monocle. Spermatogenesis being a unidirectional differentiation process, control germ cells were arranged into a linear trajectory (**Fig. 4a**). By contrast, *Dnmt3A* mutant germ cells adopted different branches, as it would be the case for a complex, multi-path differentiation process. The four SSCs subtypes contributed to the pseudotime branches (**Fig. 4b**), likely indicative of a blockage. The same trend was observed in *Dnmt3L^KO^* germ cells (**Extended data Fig. 6a,b**).

**Fig. 4.**
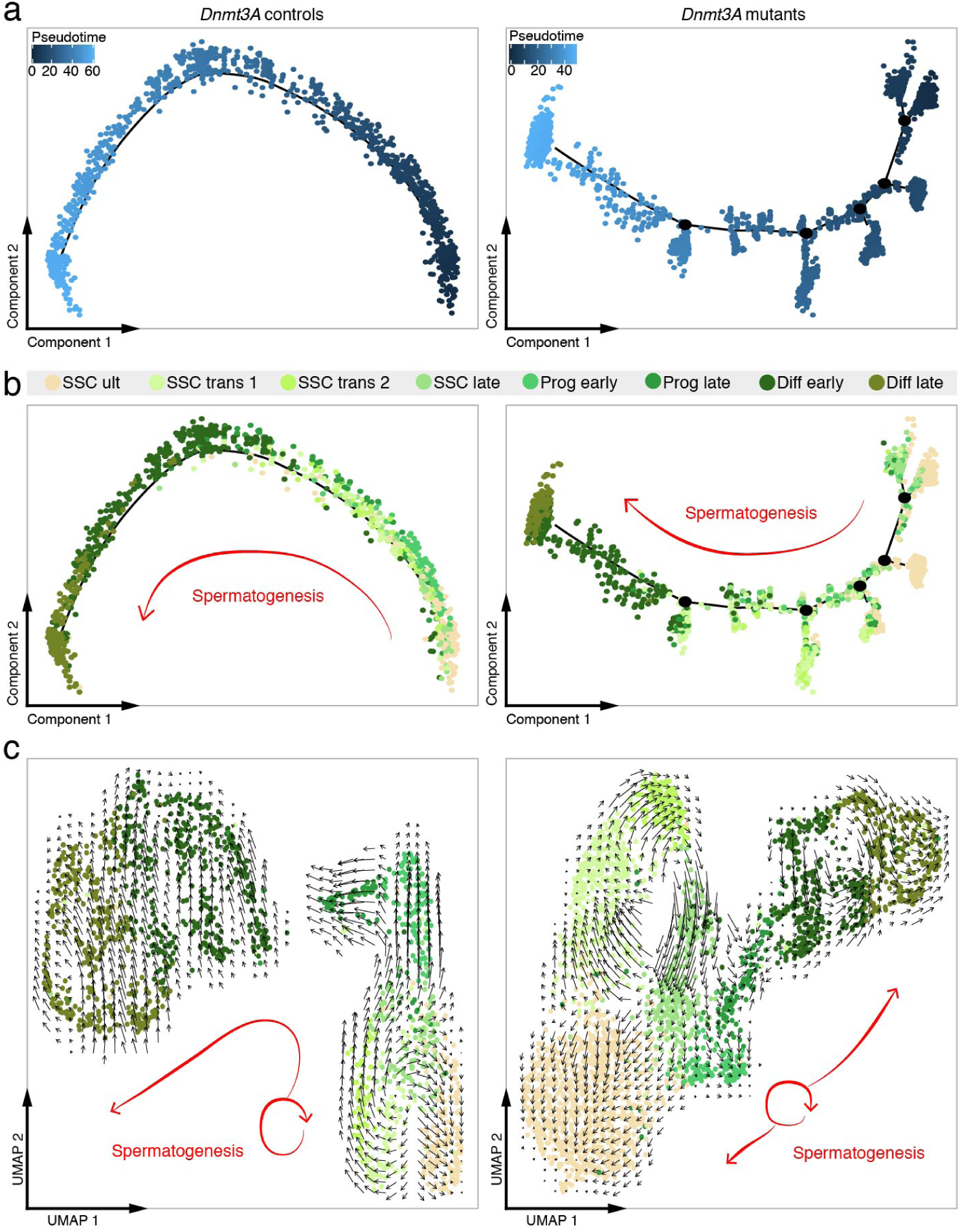
DNMT3A is essential to SSC plasticity. **a, b**, Pseudotime trajectory of germ cells from 1Odpp testes. (Left) *Dnmt3A* control germ cells (FACS-sorted *Dnmt3AWT* and *Dnmt3ACTL).* (Right) *Dnmt3A* mutant germ cells (FACS-sorted *Dnmt3A^KO^* and *Prdm1-Dnmt3N^K0^).* In (a), cells were ordered from beginning (dark blue) to the end (light blue).ln (b), cells were colored following clust,er allocation, see key above. c, Visualization of RNA velocity analysis on UMAP plot of control germ cells (Left) and *Dnmt3A* mutant germ cells (Right). Direction and size of arrows predict the future cellular state.

To further understand the dynamics of *Dnmt3A* mutant SSCs, we relied on RNA velocity analysis, which predicts the future state of a cell based on spliced versus unspliced mRNA balance^32^. With this tool, transcriptional trajectories are denoted as vectors of different amplitudes and directions. When applied to control cells, this analysis clearly illustrated SSC plasticity: SSC ultimate, transitory 1, 2 and late were all organized into a cycling process (**Fig. 4c**). Forward and backward movements between SSC subtypes likely relate to their uncommitted behaviors. In contrast, progenitors and differentiated spermatogonia were organized into a unidirectional movement, suggesting irreversible commitment towards differentiation. Strikingly, in *Dnmt3A* mutants, SSC ultimate cells no longer cycled with the other SSC subtypes. Instead, they were arranged on a unidirectional direction, opposite to spermatogenesis progression. This can be interpreted as an inability to change state and differentiate.

These findings indicate that SSCs proliferate normally but are unable to exit the stem cell pool in *Dnmt3A* mutants, which explains the spermatogenic failure. Importantly, we observed the same aberrant developmental program in *Dnmt3L* mutants: lack of DNA methylation marks established by DNMT3A prior to birth—rather than constitutive lack of DNMT3A *per se*—likely underpins the loss of SSC plasticity in *Dnmt3A* mutants.

### DNA methylation regulates SSC plasticity by limiting enhancer activity

To identify the molecular pathways that lock *Dnmt3A^KO^* SSCs into a stem cell identity, we performed bulk RNA-seq on pooled ID4-GFP^bright^ and ID4-GFP^med^ populations (ID4-GFP^bright+med^) at 10dpp, to recover ultimate SSCs and their early transitory derivatives (**Extended data Fig. 3b** and **Supplementary Table 1**). We found 726 significantly misregulated genes in *Dnmt3A^KO^* compared to WT cells (FDR < 5% and log2FC >1): 440 upregulated and 286 downregulated (**Fig. 5a** and **Supplementary Table 3**). Upregulated genes were mostly enriched in genes required for SSC homeostasis and maintenance (*Gfra1*, *Id4*, *Pax7*), while downregulated genes were involved in SSC differentiation (*Stra8*, *Kit, Sox3*) (**Fig. 5b**). Variable composition of the ID4-GFP^bright+med^ pool between *Dnmt3A^KO^* and WT could underlie the observed transcriptional changes. However, using our scRNA-seq data, we confirmed that SSC-specific markers were upregulated in individual *Dnmt3A^KO^* SSCs, from ultimate to late subtypes (**Fig. 5c**). Transcriptional features linked to SSC identity are therefore intrinsically exaggerated in *Dnmt3A^KO^* SSCs.

**Fig. 5.**
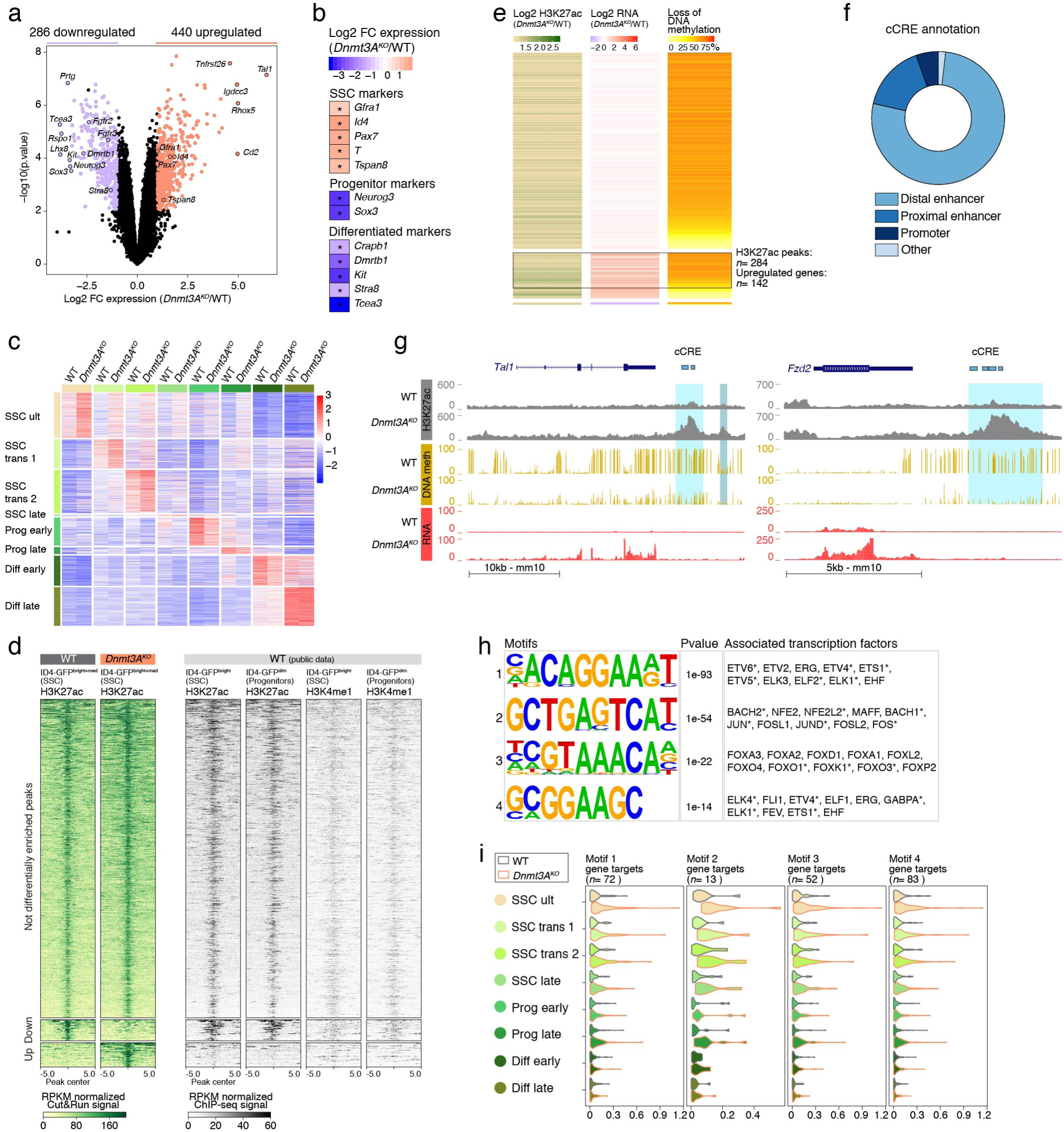
DNA methylation controls stem cell gene expression by limiting enhancer activity in SSCs. **a**, Volcano plots showing differential RNA levels in 104-eGFP^bright+med^ germ cells between *Dnmt3A^KO^* and WT (1Odpp} in log2 FC versus -log1O of the P values. Upregulated genes (440) are highlighted in red,downregulated genes (286) in purple (FDR<5% and FC>2,n= 3 biologi­ cal replicates/genotype). b, RNA-seq heatmap representation of selected germ cell type-markers that are differentially expressed between *Dnmt3N °*and WT (FIDR<5% and FC>2). **c**, scRNA-seq heatmap shows normalized expression value of markers charac­ terizing each germ cell population in corresponding *Dnmt3N°* and WT germ cell types. d, (Left) Heatmap showing levels of H3K27ac peak enrichment in RPKM-normalized Cut&Run signal from *Dnmt3A^KO^* and WT ID4-GFP^bright+med^ germ cells (10dpp). Enrichment is assessed +/- 5kb from the center of the peak. Peaks are divided into three categories: not differentially enriched *(Dnmt3N °* = WT),down-enric:hed *(Dnmt3N °* < WT), up-enriched *(Dnmt3A^KO^* > WT) (FDR<5% and FC>1). Number of biological replicates: *Dnmt3N °=* 3, WT= 4. (Right) Enrichment in H3K27ac and H3K4me1 from public ID4-eGFPbtighi and ID4-eGFPdim ChlP-seq data^36^ (RPKM-normalized signal) at H3K27ac peaks described here in *Dnmt3N °* and WT. e, Heatmap focusing on i) *Dnmt3N °-gained* H3K27ac peaks (log2 FC) compared to WT (left), ii) differential expression (log2 FC) of genes associated with H3K27ac peaks (TSS +/- 5kb from the peak) between *Dnmt3N °* and WT-from ID4-eGFP^bright+med^ RNA-seq-(center), and iii) percentage of DNA methylation loss in *Dnmt3A^KO^* versus WT on H3K27ac peak genomic location-from E18.5 WGBS-(right). Rows are ordered according to gene expression changes: top,not differentially expressed;middle,upregulated; bottom,downre­gulated. f, Pie chart of genomic annotations of ENCODE candidate cis-Regulatory Elements (cCREs) overlapping with Dnmt3A^KO^-specific H3K27ac peaks associated with >30% DNA methylation loss and upregulated genes. **g**, Genome browser representations of H3K27ac enrichment (grey), DNA methylation (yellow) and RNA expression (red) for *Dnmt3A^KO^* and WT. Regions showing Dnmt3AKO de novo H3K27ac peaks associated with >30% DNA methylation loss are shaded in light blue; regions showing de novo H3K27ac peaks at regions not controlled by DNA methylation are shaded in grey-blue.h,(Left) Enriched motifs 1 to 4 found at Dnmt3N°-specific H3K27ac peaks (n= 2,438).(Center) P values of motif 1 to 4.(Right) Transcription factors (TFs) that have the best match with the enriched motif, *indicates expression in SSCs, based on 10dpp scRNA-seq data (see Extended Fig.8d). i, Violin plots showing expression levels of TF gene targets in germ cell population subtypes discriminated by scRNA-seq for *Dnmt3A^KO^* and WT.TF gene targets were defined by association with *Dnmt3A^KO^-specific* H3K27ac peaks (TSS +/- 5kb from the peak) containing one or more of the four motifs.

We then investigated whether this enhanced SSC transcriptome was reflected at the chromatin level. We focused on two histone marks whose occupancy at promoters and enhancers is anticorrelated with DNA methylation, H3K27me3 and H3K27ac, which are themselves antagonist to each other^33–35^. In *Dnmt3A^KO^* SSCs, genome-wide reduction in DNA methylation could lead to H3K27me3 spreading and aberrant gene repression, and/or to H3K27ac gain and enhancer activation. Optimized Cleavage Under Targets and Release Using Nuclease (Cut&Run) was performed to profile these two marks in SSC-enriched ID4-GFP^bright+med^ at 10dpp (**Supplementary Table 1**). We did not observe reciprocal gain and loss in H3K27me3 and H3K27ac in *Dnmt3A^KO^* compared to WT and therefore analyzed these two marks independently (**Extended data Fig. 7a**). With regards to H3K27me3, we identified 7,139 peaks in *Dnmt3A^KO^* and WT cells, among which 15.5% (*n*= 1,108) were present in *Dnmt3A^KO^* only, and were referred to as “up-enriched” peaks (**Extended data Fig. 7b,c** and **Supplementary Table 4)**. When associated to the nearest genes (<5kb from the TSS), 33 of these genes were downregulated and conjointly lost DNA methylation in *Dnmt3A^KO^* cells (>30% compared to WT), at the same regions where H3K27me3 was gained (**Extended data Fig. 7d**). However, most *Dnmt3A^KO^*-gained H3K27me3 peaks (>75%) were intragenic (**Extended data Fig. 7e**): rather than directly repressing genes, these H3K27me3 peaks likely arose in *Dnmt3A^KO^* SSCs as a consequence of transcriptional silencing.

When considering enhancer-enriched H3K27ac marks, we counted a total of 36,431 peaks, among which 6.7% (*n*= 2,438) were present in *Dnmt3A^KO^* only (**Extended data Fig. 8a,b** and **Supplementary Table 4)**. Investigating public ChIP-seq data from ID4-GFP^bright^ and ID4-GFP^dim^ cells from prepuberal testes^36^, H3K27ac “up-enriched” peaks were neither referenced, nor decorated by H3K4me1 in one or the other cell type (**Fig. 5d**). This indicates that *Dnmt3A^KO^*-gained H3K27ac peaks in ID4-GFP^bright+med^ cells did not relate to active or primed enhancers that are normally present in ID4-GFP^bright^/SSC ultimate or ID4-GFP^dim^/progenitor cells, but were a specific feature of the *Dnmt3A^KO^* chromatin landscape.

Association of the *de novo* peaks with nearest genes revealed a total of 142 upregulated genes located in the vicinity of 284 regions that gained H3K27ac enrichment when losing DNA methylation in *Dnmt3A^KO^* germ cells (**Fig. 5e** and **Supplementary Table 4)**. This suggests that a cohort of enhancers may ectopically form in hypomethylated *Dnmt3A^KO^* SSCs. Interestingly, about half (50.7%) of the 284 *Dnmt3A^KO^*-specific and DNA methylation-restricted H3K27ac peaks overlapped with one or more ENCODE candidate *cis*-Regulatory Elements (cCREs), the vast majority being annotated as distal enhancers in other tissue types (**Fig. 5f** and **Supplementary Table 4**). This was exemplified by *Tal1*, a transcription factor involved in hematopoietic development. While not substantially expressed in normal SSCs, *Tal1* was strongly activated in *Dnmt3A^KO^* SSCs (6-fold) in association with a region that transitioned from DNA methylated to H3K27ac-occupied and that encompassed two known hematopoietic stem cell enhancers^37^ (**Fig. 5g**). Interestingly, a second H3K27ac peak was gained nearby, but in a DNA methylation-independent manner—this position was not methylated in WT SSCs—indicating a secondary recruitment, as also observed at the *Gata2* gene, another hematopoietic transcription factor (**Extended data Fig. 8c**). Genes expressed in SSCs could also further increase expression upon gaining DNA methylation-restricted H3K27ac peaks in *Dnmt3A^KO^*: this was the case for *Fzd2*, a Wnt receptor important for SSC specification and proliferation^23^ (**Fig. 5g**).

Finally, we searched for transcriptional motif signatures underlying *Dnmt3A^KO^*-specific enhancers, considering the whole set of gained H3K27ac peaks to reach enough confidence (*n*= 2,438). Using HOMER, we identified four motifs that were significantly over-represented (p < 1e^-14^) and all had potential CpG dinucleotides in their reconstructed consensus sequence. These motifs were predicted to be targeted by 35 transcription factors (TFs) that mainly belonged to ETV, ETS, ELK and FOXO families (**Fig. 5h**). Importantly, none of these TFs were significantly overexpressed in single *Dnmt3A^KO^* cells (scRNA-seq data), whether they were normally expressed in SSCs or not (**Fig. 5h** and **Extended data Fig. 8d)**. The gain of enhancers in *Dnmt3A* mutants is therefore not linked to increased TF availability but rather to increased accessibility of the underlying binding motifs, likely because of the lack of DNA methylation. Incidentally, ETV, ETS, ELK and FOXO-type TFs have been reported to be DNA methylation-sensitive and notably gain novel binding sites in DNA methylation-deficient embryonic stem cells^6, 38, 39^. A number of 116 putative gene targets were then defined by their proximity to one or more motif-containing *Dnmt3A^KO^*-specific H3K27ac peaks. These were more expressed in SSCs than in progenitors and differentiated cells in scRNA-seq data, and they were clearly upregulated in *Dnmt3A^KO^* SSC subtypes—and the most acutely in SSC ultimate—compared to WT (**Fig. 5i**). Most saliently, all but one putative gene targets belonged to the list of 142 upregulated genes directly impacted by the lack of DNMT3A-dependent DNA methylation (**Extended data Fig. 8e** and **Supplementary Table 4**).

Taken all together, these results reveal that lack of DNMT3A-dependent DNA methylation is associated with ectopic recruitment of enhancers in SSCs, which activate or enhance expression of genes that buttress SSC identity.

## Discussion

The process of *de novo* DNA methylation is an early event of spermatogenesis, occurring prior to birth, before SSC emergence and meiosis. We show here that DNMT3A and DNMT3C jointly shape the fetal male germline methylome with perfect complementarity and striking lack of redundancy in the genomic sequences they target, and most importantly, in the spermatogenetic phase they control. While it was known that DNMT3C is crucial for meiosis, by selectively methylating and silencing retrotransposons^12^, we found that DNMT3A-dependent DNA methylation is widespread and regulates SSC homeostasis. More precisely, the hypomethylated genomic landscape of *Dnmt3A* mutant SSCs results in appearance of ectopic enhancers that reinforce the robustness of the stem cell gene expression program, which very likely locks SSCs into self-renewal and prevents commitment to differentiation. Collectively, we provide evidence for a novel reproductive function for DNA methylation, beside meiosis protection: the programming of life-long spermatogenesis fueling (**Fig. 6**).

**Fig. 6.**
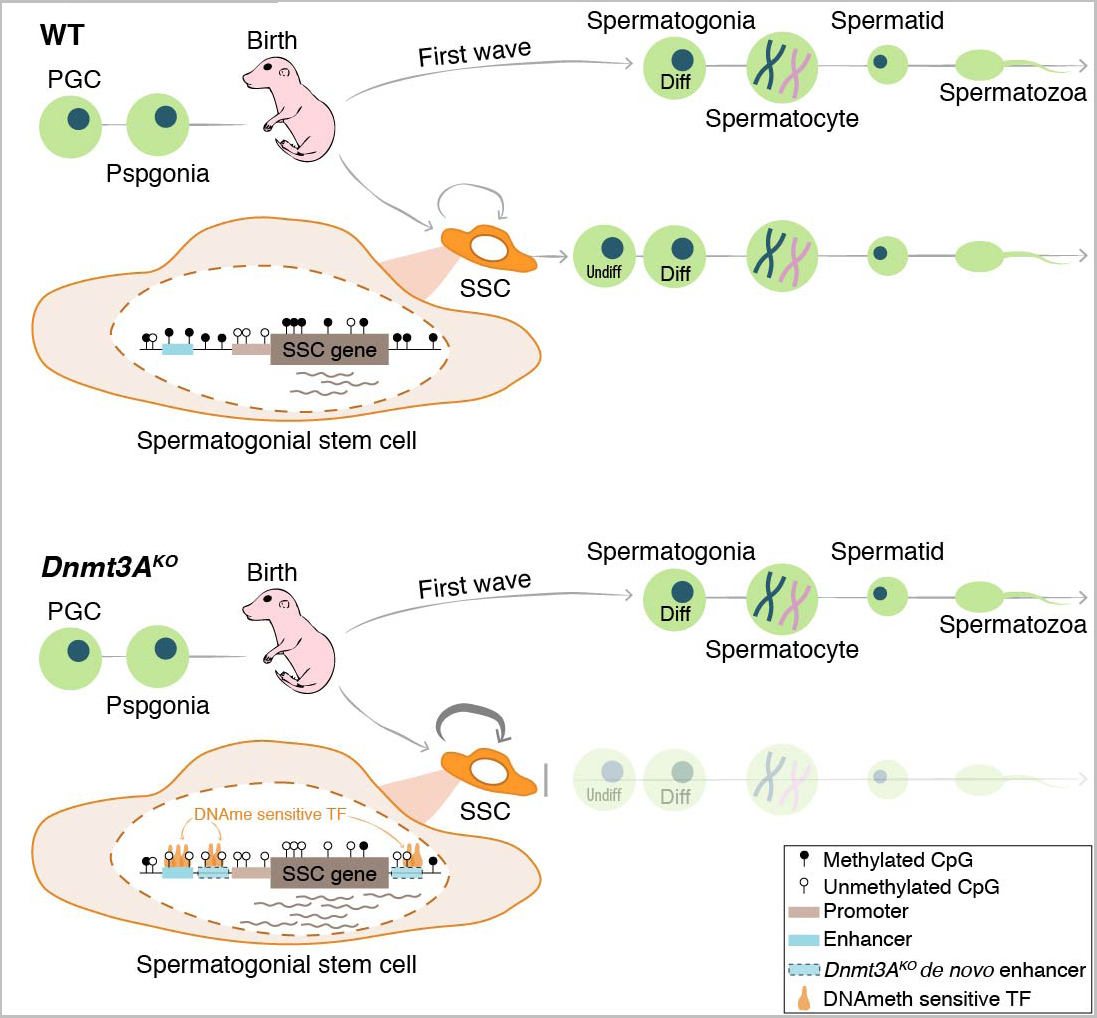
Model of DNMT3A-dependent DNA methyla­ tion in spermatogenesis. Top, Developmental options of fetal prospermatogonia (Pspgonia) at birth in WT males: i) directly transition to differentiated spermatogo­nia (Oiff) to initiate the first wave of spermatogenesis, or ii) form the foundational SSC pool, which must balance self-renewability and differentiation to sustain sperma­ togenesis across male reproductive lifespan. Bottom, lifelong reproductive consequences of failure to establish DNA methylation in *Dnmt3N °* Pspgonia: i) the first wave of SSC-independent spermatogenesis can proceed, however ii) once prospermatogonia develop into SSCs, these can only self-renew and are unable to differentiate, preventing following waves of spermatoge nesis. The hypomethylated genome uncovers ectopic enhancers (dashed lines) that become accessible to methyl-sensi­ tive TFs (orange) and enforce irreversible expression of SSC-promoting genes.

This study resolves several questions and controversies regarding DNMT3A role in spermatogenesis. First, conflicting reports existed as to whether *Dnmt3A^KO^* germ cells can undergo meiosis or not^11, 15^. We demonstrate that *Dnmt3A* mutant males complete meiosis during the first round of spermatogenesis—which bypasses the SSC state—and can generate haploid cells, including spermatozoa. However, they produce post-meiotic cells only once in their life, after which their germ cell contingent exclusively consists in differentiation-deficient SSCs. Second, we definitely exclude a role for DNMT3A in retrotransposon control during spermatogenesis; DNMT3C is solely dedicated to this pathway in mice. Coincidently, we prove that meiotic recombination remarkably tolerates low genomic methylation levels, provided that retrotransposon promoters are methylated. Finally, NSD1-dependent H3K36me2 deposition was recently shown to drive *de novo* DNA methylation of euchromatic regions in fetal male germ cells^40^. We reveal that *Nsd1^KO^* and *Dnmt3A^KO^* prospermatogonia share a genome-wide pattern of hypomethylation that excludes young retrotransposon promoters, highlighting that DNMT3A is the enzyme recruited to H3K36me2-marked regions, likely through its PWWP reading domain. However, despite convergent DNA methylation defects, we do not expect a convergent phenotype in *Dnmt3A* and *Nsd1* mutants.

Accumulation of H3K27me3 leads to specific gene dysregulation in *Nsd1^KO^* germ cells, a feature that was not observed in *Dnmt3L^KO^* germ cells^40^, and which we did not observe in *Dnmt3A^KO^* SSCs. Moreover, *Nsd1* mutants exhibit early spermatogonia loss^40^: this contrasts with the expanding SSC phenotype we uncovered in *Dnmt3A* and *Dnmt3L* mutants. This discrepancy underscores the complexity of the epigenetic interplay that regulates male germ cell development.

The spermatogenetic failure of *Dnmt3A* mutant males is quite unique among other SSC-related infertility phenotypes. Disrupted SSC homeostasis and subsequent spermatogenetic decline have been previously linked to defects in generating the foundational SSC compartment, to impaired self-renewal and precocious exhaustion of the SSC pool, or to defective differentiation priming^41, 42^. Remarkably, *Dnmt3A^KO^* SSCs lose their plasticity by continuously self-renewing and ignoring the differentiation route. Through chromatin profiling combined with bulk and single-cell transcriptomics, we were able to provide detailed mechanistic insights into this phenotype: SSC expansion is not linked to hyperproliferation but rather to an intrinsic exacerbation of a stem cell program. A cohort of ectopic H3K27ac-enriched enhancers arise in *Dnmt3A^KO^* SSCs, and this is linked to upregulation of nearby SSC-specific genes but also, notably, to the activation of stem cell genes that are normally relevant to other tissue-specific stem cells, such as hematopoietic stem cells. We found that these enhancers are largely methylated by DNMT3A during fetal development, prior to the formation of the SSC pool, and most importantly, they possess binding motifs for methyl-sensitive TFs, such as the ETV- or ETS-type TF families^38, 43^.

Interestingly, in proximity to these DNA methylation-restricted enhancers that gained H3K27ac in *Dnmt3A^KO^* SSCs, we frequently observed additional enhancers that emerged at regions that were not controlled by DNA methylation, indicating secondary reinforcement of the SSC gene program through additive and synergistic TF binding. We propose that DNMT3A-dependent methylation, assisted by DNMT3L in fetal germ cells, programs SSC dual capacity to both self-renew and differentiate after birth, by limiting the accessibility of stem cell-related enhancers to methyl-sensitive TFs that are abundantly present in SSCs.

The essential role of DNMT3A has been previously described in other stem cell systems, although mechanisms at play may be different and/or not fully resolved. In neural stem cells (NSCs), DNMT3A-dependent DNA methylation does not limit but rather activate enhancers linked to neurogenesis, by antagonizing H3K27me3^44^. Moreover, *Dnmt3A^KO^* NSCs are normal but undergo skewed differentiation towards a glial rather than neuronal fate. In hematopoietic stem cells (HSCs), *Dnmt3A* deficiency is also not impactful in steady-state-conditions but skewed trajectory occurs upon differentiation, in association with increased accessibility of CpG-rich TF binding motifs linked to the myeloid lineage^45, 46^.

Interestingly, when challenged by serial transplantation, the *Dnmt3A^KO^* HSC compartment increasingly expands towards unlimited self-replication, while differentiation potential inversely declines^47, 48^. This is reminiscent to the effect we found in *Dnmt3A^KO^* SSCs, suggesting that the molecular mechanisms we evidenced could also apply to transplanted *Dnmt3A^KO^* HSCs. Overall, differences in DNMT3A role in various stem cell-supplied systems may be attributable to distinct physiology and epigenetic dynamics. More precisely, spermatogenesis is a unidirectional process, while hematopoietic or neural differentiation are multilineage paths. Then, DNA methylation is dramatically reduced in *Dnmt3A^KO^* SSCs (by 3 to 4 fold compared to WT), while changes are relatively sparse and focal in *Dnmt3A^KO^* HSCs or NSCs. As demonstrated here, DNMT3A is the main *de novo* methylation enzyme in embryonic germ cells, while its activity is less important in embryonic somatic cells, whose genome mostly acquires DNA methylation in a DNMT3B-dependent manner^49^.

To conclude, these results provide novel and significant advances towards understanding the regulation of stem cell function by DNMT3A and the relationship between male infertility and epigenetic anomalies.

## Author contributions

M.D. and D.B. designed and conceived the study. Most experiments were performed and analyzed by M.D. M.A. assisted in the histology, immunofluorescence and microscopy. J.B. performed the WGBS experiment. L.B. and M.A. assisted in the Cut&Run experiment. L.G.B and S.L. performed NGS experiments and supervised generation of single cell and NGS data. A.T. performed the bioinformatic analyses. M.W. provided the *Dnmt3A* mouse strain. D.B. and M.D. interpreted data and wrote the paper. All authors reviewed and approved the final manuscript.

## Methods

### Mice

Mice were hosted on a 12h light/12h dark cycle with free access to food and water in the pathogen-free Animal Care Facility of the Institut Curie (agreement C 75-05-18). All experimentations were approved by the Institut Curie Animal Care and Use Committee and adhered to European and national regulation for the protection of vertebrate animals used for experimental and other scientific purposes (directives 86/609 and 2010/63). For tissue and embryo collection, mice were euthanized by cervical dislocation.

Mouse strains, all bred onto C57Bl6/J background, were previously described: *Dnmt3A^2lox^* ^11^, *Dnmt3L^KO^* ^10^, *Dnmt3C^KO^* ^12^, *Prdm1-Cre* ^21^ (gift of A. Clark, UCLA), *Id4-eGFP* ^24^ (gift of J. Oatley, Washington State University) and *Oct4-eGFP* (Jackson Laboratories, stock no. 008214)^17^ (gift of A. Clark, UCLA). The *Dnmt3A^KO^* strain was obtained by constitutive defloxing of the *Dnmt3A^2lox/2lox^* strain using the *Zp3-Cre* line (Jackson Laboratories, stock no. 003394)^50^. Germ-cell conditional *Dnmt3A* mutants were obtained by crossing *Dnmt3A^KO/WT^*; *Prdm1-Cre^Tg/0^* males with *Dnmt3A^2lox/2lox^* females. The mutant genotype of interest was *Dnmt3A^KO/2Lox^*; *Prdm1-Cre^Tg/0^* (referred to as *Prdm1-Dnmt3A^cKO^* in figures and text); control genotypes were *Dnmt3A^2lox/WT^*; *Prdm1-Cre^Tg/0^*, *Dnmt3A^KO/WT^*; *Prdm1-Cre^Tg/0^*, *Dnmt3A^2lox/KO^*; *Prdm1-Cre^0/0^*, or *Dnmt3A^2lox/WT^*; *Prdm1-Cre^0/0^*, as indicated in legends of each experiment (referred to as *Dnmt3A* controls).

Prenatal time points were obtained by following timed pregnancies where the following day post-coitum was considered as embryonic day E0.5. Postnatal time points were measured starting from birth, considering the first postnatal day as 1 day post-partum (1dpp).

### RT-qPCR and RT-PCR

Testes (one testis per biological replicate) were homogenized in Trizol (Life Technologies) using TissueLyser LT (Qiagen). Total RNA was extracted according to the manufacturer’s recommendations, and DNase-treated with the Qiagen RNase-Free DNase Set. RNA was quantified using Qubit Fluorometric Quantitation (Life Technologies), and checked for integrity using the TapeStation (Agilent). For RT-PCR and RT-qPCR, 2µg of RNA was reverse transcribed using random priming with SuperScript III (Life Technologies). RT-PCR and qRT-PCR primers are listed in **Supplementary Table 5**.

Quantitative PCR (RT-qPCR) was conducted using Power SYBR Green PCR Master Mix (Life Technologies) on a ViiA7 Real-Time PCR System (Applied Biosystems). Data were normalized to *beta-actin* using the delta-delta Ct approach. Minus RT controls were included for each collected data point.

PCR (RT-PCR) was conducted using GoTaq DNA polymerase (Promega). PCR programs were the following, *Sp10*: 30 cycles (94°C for 30 sec; 60°C for 30 sec; 72°C for 1 min) and *Tp2*: 30 cycles (94°C for 30 sec; 58°C for 30 sec; 72°C for 30 sec). Minus RT controls were included for each collected data point.

### Histological sections

Testes were dissected without disrupting the tunica albuginea, fixed over-night (ON) in fresh Davidson’s-modified 4M formaldehyde fixative (4M formaldehyde, 15% ethanol, 5% glacial acetic acid), washed (2×10min and 1×30min) in 1XPBS with 0.25M Tris-HCl pH7.2 and then stored at 4°C in 1xPBS-70% ethanol with 0.25M Tris-HCl pH7.2. Alternatively, testes were fixed ON in Bouin’s fixative solution (Sigma), washed and stored in 70% ethanol at 4°C.

Testes were then paraffin-embedded, sectioned (8 µm) and stained with periodic-acid-Schiff (PAS) using standard protocols. Slides were imaged on a Leica upright epifluorescence microscope using color camera. For quantification of tubule types (**Fig. 2a** and **Extended data Fig. 2e**), tubules from one testis section per animal were counted. This represents above 100 tubules for all ages and genotypes, except at 6 months for *Dnmt3A^KO^* and *Dnmt3L^KO^* genotypes (due to severe testis size reduction), where around 60 tubules were counted per section.

### Immunofluorescence microscopy

For immunofluorescence on testis sections, testes were collected, fixed at 4°C in 4% paraformaldehyde (2dpp: 1h; 10dpp: 2h; 25dpp: 2h30; adult: ON), washed 2×10min and 1×30min in 1XPBS with 0.25M Tris-HCl pH7.2 and 3×5min in 1XPBS. Then testes were soaked in 15% and 30% sucrose solution in 1XPBS with 50mM Tris-HCl pH7.4 for two consecutive ON incubations, embedded in OCT medium (Tissue-Tek) and frozen at -80°C. Frozen sections were cut (6-10 μm) and spotted onto positively-charged slides (Superfrost Plus slides-Thermo Fisher Scientific) and stored at -20°C prior to use. For immunofluorescence detection, slides were thawed at room temperature (RT), washed in 1XPBS, and then blocked and permeabilized for 1h at RT in blocking buffer (10% donkey serum, 3% BSA and 0.2% Triton in 1XPBS). Sections were incubated ON at 4°C with primary antibody diluted in blocking buffer, followed by washes (3×5min) with 1XPBS supplemented with 0.1% Tween-20, and then incubated for 1-3h at RT with Alexa Fluor-conjugated secondary antibodies diluted in blocking buffer. After final washes, sections were mounted in DTG (Glycerol, DABCO 2.5%, 50mM Tris pH8.6) with DAPI. Images were obtained using an inverted laser-scanning confocal microscope LSM 700 or 900 (Zeiss).

Each staining was performed on three different testis sections from at least three different animals. Primary antibodies were omitted as a negative control. For quantification, the number of cells were normalized by the mean of the tubule surface.

For immunofluorescence on meiotic spreads^51^, testes (16 to 25dpp) were collected, seminiferous tubules were released from the tunica albuginea, rinsed in 1XPBS and incubated for 1h in hypotonic buffer (30mM Tris, 50mM sucrose, 17mM trisodium citrate dihydrate, 5mM EDTA, 0.5mM DTT, pH 8.2) containing 0.5mM protease inhibitors (Roche). Seminiferous tubules were then placed in a drop of sucrose solution (100mM sucrose pH 8.2) and tubules were dissociated until single cell suspension. Cells were slowly dispersed on slides pre-covered of a fresh fixative solution (2% formaldehyde, 0.05% triton X-100, 0.02% SDS). Slides were air-dried, washed (3×5min) in 0.2% Photo-Flo (Kodak) and stored at -80°C until use. Immunofluorescence detection was performed as described above, except that slides were blocked for 20min at RT and incubated with primary and secondary antibody for 1h at RT. Slides were imaged on an upright epifluorescence microscope. Antibody references and working dilutions are available in **Supplementary Table 6**.

### Cell sorting and analysis by fluorescent activated cell sorting (FACS)

Testes were dissected from E18.5 or 10dpp males, decapsulated and transferred in low-binding Eppendorf tube containing 100-150µL collagenase solution in HBSS (Collagenase type IV (Gibco), 2X AAs (Gibco), 2X Na-pyruvate (Gibco), 25mM HEPES-KOH pH7.5). Dissociation was achieved at 37°C for 5-7min by gently flicking the tube. Then 400-600µL of TrypLE Express (Gibco) was added to the cell suspension and incubated for 5-7min at 37°C. Single-cell suspension was obtained by up-and-down pipetting. TrypLE Express was quenched with 140-210µL fetal bovine serum (FBS) and the cell suspension was centrifuged at 600g/4min/4°C, washed with FACS buffer (1XPBS, 2mM EDTA, 1%BSA) and filtered through a 35 μm strainer.

For sorting prospermatogonia or spermatogonial stem cells (SSCs), *Oct4-eGFP* and *Id4-eGFP* transgenic mice were used, respectively. Washed single-cell suspension was centrifuged and resuspended in FACS buffer supplemented with DAPI (2ng/ml), in order to discriminate dead cells. Cell sorting was performed on a BD FACSaria II (BD Biosciences) using 100 μm nozzle. Prospermatogonia were isolated by gating DAPI-neg, OCT4-GFP-pos cells. SSCs were isolated by gating DAPI-neg, ID4-GFP-pos cells. ID4-GFP positive cells were further separated into three size-matched compartments (bright, medium and low); only ID4-GFP^bright^ and ID4-GFP^med^ were sorted and considered as SSCs (ID4-GFP^bright+med^)(see gates **Extended data Fig. 3c**).

For spermatogonia sorting and analysis, single-cell suspensions were incubated 20min on ice in 500µL FACS buffer containing 3.5µg/mL of rat anti-mouse EpCAM antibody conjugated with Alexafluor 647 (Biolegend) and 8µg/mL mouse anti-β2-Microglobulin monoclonal antibody conjugated with PE (Santa Cruz). After antibody labeling, cells were washed in FACS buffer and resuspended in DAPI-FACS buffer. Compensation controls were applied for each experiment. Cell sorting and data acquisition were performed on FACSaria II (BD Biosciences) using 100 μm nozzle. Spermatogonia were isolated by gating DAPI-neg, EpCAM-pos, β2M-neg cells. Further analyses were performed with FlowJo software.

### DNA methylation analyses

WGBS was performed on prospermatogonia sorted by FACS (with the *Oct4-eGFP* transgenic line) using direct post-bisulfite adaptor tagging (PBAT)^18^. Prospermatogonia of three embryos from each genotype were pooled for genomic DNA extraction using the QIAamp DNA Micro Kit (Qiagen) and spiked-in with 0.3% of unmethylated Lambda DNA (Promega). Bisulfite conversion was performed using EZ DNA Methylation-Gold Kit (Zymo Research) and the whole converted DNA was used as input for library preparation using the Accel-NGS Methyl-Seq DNA Library Kit (Swift Biosciences) according to the recommended protocol and indexing PCR using 2X KAPA HiFi HotStart Uracil+ ReadyMix (KAPA Biosystems). Indexed, cleaned-up libraries were analyzed and quantified on a 2200 Tapestation instrument using a D5000 screen tape (Agilent). Library pools were sequenced on an Illumina NovaSeq in 100bp paired-end (PE) reads run.

Targeted analysis of DNA methylation at retrotransposon promoters (IAPEz, L1MdA and L1MdTf) was performed on FACS-sorted spermatogonia (gating DAPI-neg, EpCAM-pos, β2M-neg cells) at 10dpp. DNA was extracted by incubating one volume of lysis buffer (200mM Tris-HCl pH8, 10mM EDTA, 0.4% SDS, 400mM NaCl, 80µg/mL linear polyacrylamide, 0.4mg/mL proteinase K) at 55°C for 2-4h, precipitated with isopropanol and cleaned with 70% ethanol. DNA was then bisulfite-converted by using EZ DNA Methylation-Lightning^TM^ Kit (Zymo Research). CpG methylation was quantified on at least three individual animals of each genotype by pyrosequencing on a PyroMark Q48 (Qiagen) using the Q48 Software. Primers are available in **Supplementary Table 5**.

### RNA sequencing

For bulk RNA-seq of whole testes, testes were collected from *Dnmt3A^KO^* and WT littermate animals at 19dpp and 25dpp (*n*=2 per genotype and age) and homogenized in Trizol (Life Technologies) using TissueLyser LT (Qiagen). Total RNA was extracted according to the manufacturer’s recommendations, and DNase-treated with the RNase-Free DNase Set (Qiagen). RNA was quantified using Qubit Fluorometric Quantitation (Life Technologies), and checked for integrity using TapeStation (Agilent). RNA-seq libraries were performed with TruSeq Stranded mRNA Kit (Illumina) and sequenced in 100bp PE reads run on NovaSeq 6000 (Illumina). Published data for *Dnmt3C* mutants (*Dnmt3C^IAP/IAP^*) at 20dpp were including in the analysis^12^.

For bulk RNA-seq of SSCs, ID4-GFP^brigth+med^ germ cells from 10dpp mice were FACS-sorted as described above and 5,000-10,000 cells were collected for each animal (*n*= 3 per genotype) in low binding Eppendorf tubes pre-filled with 100µL of Extraction Buffer from Arcturus PicoPure RNA isolation Kit (Applied Biosystems). Samples were frozen and stored at -80°C for later use. RNA extraction was performed following detailed protocol of the same kit and DNase-treated with RNase-Free DNase Set (Qiagen). RNA concentration and quality were assessed by Bioanalyzer RNA 6000 Pico Assay (Agilent). RNA-seq libraries were performed with SMARTer® Stranded Total RNA-Seq Kit v2 (Takara - rRNA depletion) and sequenced in 100bp PE reads run on NovaSeq 6000 (Illumina), with three biological replicates per genotype.

For single cell RNA-seq (scRNA-seq), germ cells were enriched by FACS from 10dpp mice as described above (gating DAPI-neg, EpCAM-pos). Cell suspensions were loaded into Chromium Single Cell 3’ Kit (V2 or V3) and used to isolate single-cell in Gel Bead-in-Emulsion (GEMs) using the Chromium controller (10X Genomics). In all cases, cells were loaded in the proper concentration on the instrument with the expectation of collecting up to 6,000 GEMs containing cells per samples. Single-cell libraries were performed according to manufacturer’s recommendations of Chromium (10X Genomics) and sequenced in 26-91bp paired-end reads run for V2 samples (*Dnmt3L^WT^* and *Dnmt3L^KO^*) and 28-91bp paired-end reads run for V3 samples (all other samples) on NovaSeq 6000 (Illumina).

### Cut&Run

The Cut&Run protocol was modified from Skene et al. (2017)^52^. In brief, 20µL of Concanavalin A beads (Polysciences) per sample were resuspended in 1mL of Binding Buffer (20mM HEPES-KOH, 10mM KCl, 1mM CaCl2, 1mM MnCl2). Beads were washed twice in 1mL Binding Buffer and resuspended in 20µL of Binding Buffer/ sample.

ID4-eGFP^bright+med^ germ cells (between 5,000-10,000 cells) were sorted by FACS into 1X PBS (∼50µL), as described before. Samples were split into aliquots according to the number of antibodies profile required and 1mL of Wash buffer (20mM HEPES-KOH, 150mM NaCl, 0.5mM spermidine (Sigma) and 1X Complete™ EDTA-free protease inhibitor cocktail (Roche) was gently added to the cell solution and 20µL of pre-washed beads. Cells and beads were incubated for 10min at RT on a rotating wheel. Cells were then collected on magnetic beads, the supernatant was discarded and cells were resuspended in 400µl of Antibody buffer (Wash buffer with 2mM EDTA, 0.05% digitonin (Millipore) and 1:200 antibody). Cells were incubated with antibodies for 1h at RT on a rotating wheel and washed twice with 1ml of Dig wash buffer (Wash buffer with 0.05% digitonin). Antibody references are available in **Supplementary Table 6**.

Samples were then incubated with 1:400 ProteinA-MNase fusion protein (produced by the Recombination Protein Platform of the Institut Curie, 0.785 mg/mL) for 15 min at RT followed by two washes with 1ml Dig wash buffer. Cells were then resuspended in 150µl Dig wash buffer and cooled down on ice for 5min before addition of CaCl2 to a final concentration of 2mM. Targeted digestion was performed for 30 min on ice, reaction was stopped by adding 150µL of 2X STOP buffer (340mM NaCl, 20mM EDTA, 4mM EGTA, 0.02% Digitonin, 0.125mg/mL RNase A, 0.25mg/mL Glycogen). Samples were then incubated at 37°C for 20min to release cleaved chromatin fragments into the supernatant. After centrifugation at 15,000rpm for 5min, supernatants were transferred to new low-binding tubes. Following addition of 0.1% SDS and 0.17mg/mL Proteinase K, samples were mixed by inversion and incubated at 70° C for 30 minutes. DNA was purified using phenol/chloroform followed by chloroform extraction and precipitated with 10µg of glycogen and 3 volumes of 100% ethanol for at least 20 minutes on ice. DNA was pelleted at 14,000 rpm at 4°C for 20 minutes. The DNA pellet was washed in 85% ethanol, spin down and resuspended in 40µL low Tris-EDTA (10mM Tris, 0.1mM EDTA) after complete evaporation of the ethanol.

Library preparation was performed according to manufacturer’s instructions (Accel-NGS 2S DNA library kit, Swift biosciences) with a modified library amplification program: 98°C for 45sec, (98°C for 10sec, 60°C for 15sec, 68°C for 1min)x15 cycles, hold at 4°C. Average library size was tested on Agilent 4200 Tapestation using a DNA5000 screentape and quantification was performed on Invitrogen QUBIT 4 using high sensitivity DNA kit. Cut&Run libraries were sequenced on a NovaSeq (Illumina) using PE 100bp run, with four biological replicates for *Dnmt3A^WT^* and three for *Dnmt3A^KO^*.

### WGBS analysis

Adapters were trimmed using Atropos (v1.1.16). Trimmed reads were cleaned by removal of 5bp in 5’ end of read1 and 12bp in 5’ end of read2 using Cutadapt v1.12. Reads shorter than 15bp were discarded. Cleaned reads were aligned onto the Mouse Reference Genome (GRCm38/mm10) using Bismark v0.18.2 with Bowtie2-2.2.9 allowing one mismatch in a seed alignment. Only reads mapping uniquely to the genome were kept, and methylation calls were extracted after duplicate removal by considering only CpG dinucleotides covered by a minimum of five reads. CpG methylation levels over different genomic compartments were calculated by extracting methylation calls with positional overlap with coordinates for Gencode vM16 gene annotations (“Intragenic”) and the RepeatMasker database (“Transposons”). CpG islands (“CGIs”) were defined previously^54^. “Intergenic” partitions were defined as genomic regions that did not overlap with CGIs, intragenic regions or transposable elements. Differentially Methylated Region (DMR) calling was performed using the bioconductor package DSS^55^ with the following parameters: a CpG methylation level difference of at least 25%, at least five CpGs called, minimum length of 200 bp, and at least 500 bp between two DMRs. CpG methylation data metaplots were processed using deepTools v2.5.3; only individual element annotations with size greater than 5 kb were analyzed for L1s, and greater than 500bp for MERVL and IAPEz (to account for solo LTR elements).

### RNA-seq analysis

Adapters were trimmed using Atropos v1.1.16. Paired-end read alignment was performed onto the Mouse Reference Genome (mm10) with STAR (v2.7.0a) reporting randomly one position and allowing 6% mismatches (*--outFilterMultimapNmax 5000 --outSAMmultNmax 1 --outFilterMismatchNmax 999 --outFilterMismatchNoverLmax 0.06*). Repeat annotation was downloaded from RepeatMasker (http://www.repeatmasker.org/). To reconstruct full-length LTR copies, we used the perl tool “*One code to find them all”.* Reconstructed transposon annotation and basic gene annotation from GENCODE v18 were merged and used as input for quantification with FeatureCounts v1.5.1. Differential expression analysis was performed using edgeR’s normalisation combined with voom transformation from limma R package. P-values were computed using limma and adjusted with the Benjamini-Hochberg correction. Genes and transposon families were declared as differentially expressed if FDR < 5% and log2FC >1.

### Single-cell RNA-seq analysis

Raw count matrices were generated using Cell Ranger v3.0.2 and were imported to Seurat v3.1.1. To filter out low-quality cells, cells with at least 1000 genes detected and less than 15% of mitochondrial reads were kept for the following analyses. Log-normalization procedure (*NormalizeData* function) and detection of variable genes (*FindVariableFeatures* function) were performed for each sample independently. Samples were integrated by identifying anchors (*FindIntegrationAnchors and IntegrateData* functions) with 30 dimensions. Integrated data were scaled. The non-linear dimensional reduction technique UMAP was run with the 30 principal components stored after PCA analysis on the integrated data. Clusters were defined using a resolution parameter of 0.8.

Marker genes for each cluster were determined using edgeR package for the normalization and limma package for differential expression analysis if FDR<5% and log2 Fold-change > 0.25. Cell clusters were defined as germ cell clusters if they were *Ddx4* positive (universal germ cell marker) and *Gata4* negative (general somatic marker)^57^. Some clusters had a poor-quality control with low number of UMIs and detected genes, or barely no marker gene defined or too much mitochondrial genes. These clusters were excluded from the following analyses. A second round of cell selection was performed using the same method. The final analyses were done on 6,638 retained germ cells.

After identification of germ cell clusters, raw count matrices without testicular somatic cells were re-analyzed with Seurat using the same procedure as before for normalization, sample integration, clustering (resolution parameter of 0.3) and detection of markers. The average expression (per germ cell cluster) of highly variable genes (*n*=2,000 genes) was used as input to calculate Spearman correlation between germ cell clusters.

Cell cycle analysis was performed using the *CellCycleScoring* function in Seurat. Genes specific to S and G2M phases were defined using the following criteria: expressed in spermatogonia and specific to one of the two phases based on a cell-cycle gene expression analysis^30^.

Raw count matrices without testicular somatic cells were imported to Monocle2 (v2.12.0) and only expressed genes above threshold (0.1) were used for analyses. When two samples were included together in a pseudotime analysis, sample batch effects were removed via a model formula^58^.

Velocyto (v0.17.16) was used to processed raw data to count spliced and unspliced reads for each gene and generated a loom file for each sample. Loom files were imported in R (v3.6.0) using SeuratWrappers (v0.1.0) R package. RNA velocity was estimated using *RunVelocity* function and Velocyto.R package (v0.6) (using gene-relative model with k=20 cell kNN pooling and using top/bottom 2% quantiles for gamma fit). The RNA velocity map was projected onto the UMAP plot with a neighborhood size of *n*= 200 cells^32^. RNA velocity could not be performed on *Dnmt3L^KO^*, due to insufficient germ cells (1,031 WT and 1,182 *Dnmt3L^KO^*) and the use of anterior version of the scRNA-seq kit (10X Genomics V2).

### Cut&Run analysis

Paired-end reads were trimmed using Trim Galore v0.4.4. The alignment was performed onto a concatenated genome using the Mouse Reference Genome (mm10) and the *Escherichia coli* genome (str. K-12 substr. MG1655, Genbank: NC_000913) with STAR (v2.7.0e) reporting randomly one position, allowing 4% of mismatches (*--outFilterMultimapNmax 5000 --outSAMmultNmax 1 --outFilterMismatchNmax 999 --outFilterMismatchNoverLmax 0.04 --alignIntronMax 1 --alignMatesGapMax 2000*). PCR duplicates were removed using Picard v2.6.0 (http://broadinstitute.github.io/picard/). Peaks were called using MACS2 v2.1.1 using IgG sample as control. The broad option was used for

H3K27me3 samples whereas narrow peaks were detected for H3K27ac samples. Heatmap using peak centers as windows was performed using Deeptools v2.5.3. *De novo* motifs were called using HOMER v4.11 with the HOCOMOCO mouse motif database v11 as known motifs. Genomic coordinates of ENCODE candidate cis-Regulatory Elements (cCREs) were downloaded from the SCREEN website (https://screen.encodeproject.org/), and intersected with 284 *Dnmt3A^KO^*-gained H3K27ac peaks (associated with nearby gene upregulation and loss of DNA methylation in *Dnmt3A^KO^* SSCs) using bedTools (v2.27.1). Only peaks that fully overlapped with cCRE regions were considered.

## Acknowledgements

We are grateful for support and feedback from members of the Bourc’his lab, to N. Fayaubost for animal care, M. Greenberg for critical reading of the manuscript and N. Servant for bioinformatic assistance. We thank A. Clark for the *Prdm1*-Cre and *Oct4*-eGFP mice, J. Oatley for the *Id4*-GFP mouse and K. Laband and J. Dumont for the anti-SCP3 antibody. We acknowledge the ICGex NGS platform of Institut Curie (supported by grants ANR-10-EQPX-03, Equipex and ANR-10-INBS-09-08, France Génomique)- and the Cell and Tissue Imaging Platform-PICT-IBiSA (member of France-Bioimaging, ANR-10-INBS-04) of the Genetics and Developmental Biology Dpt (UMR3215/U934) of Institut Curie. The laboratory of D.B. is part of the LABEX DEEP (ANR-11-LABX-0044, ANR-10-IDEX-0001-02). This work was supported by a grant from the Agence Nationale pour la Recherche (ANR-17-CE12-00013-01), the Fondation Bettencourt Schueller and the Fondation pour la Recherche Médicale (FRM Team Label). M.D. was supported by PhD fellowships from Région Ile-de-France and Fondation pour la Recherche Médicale. L.B. is the recipient of a PhD Boehringer Ingelheim Fonds fellowship.

---

**Extended data Fig. 1.**
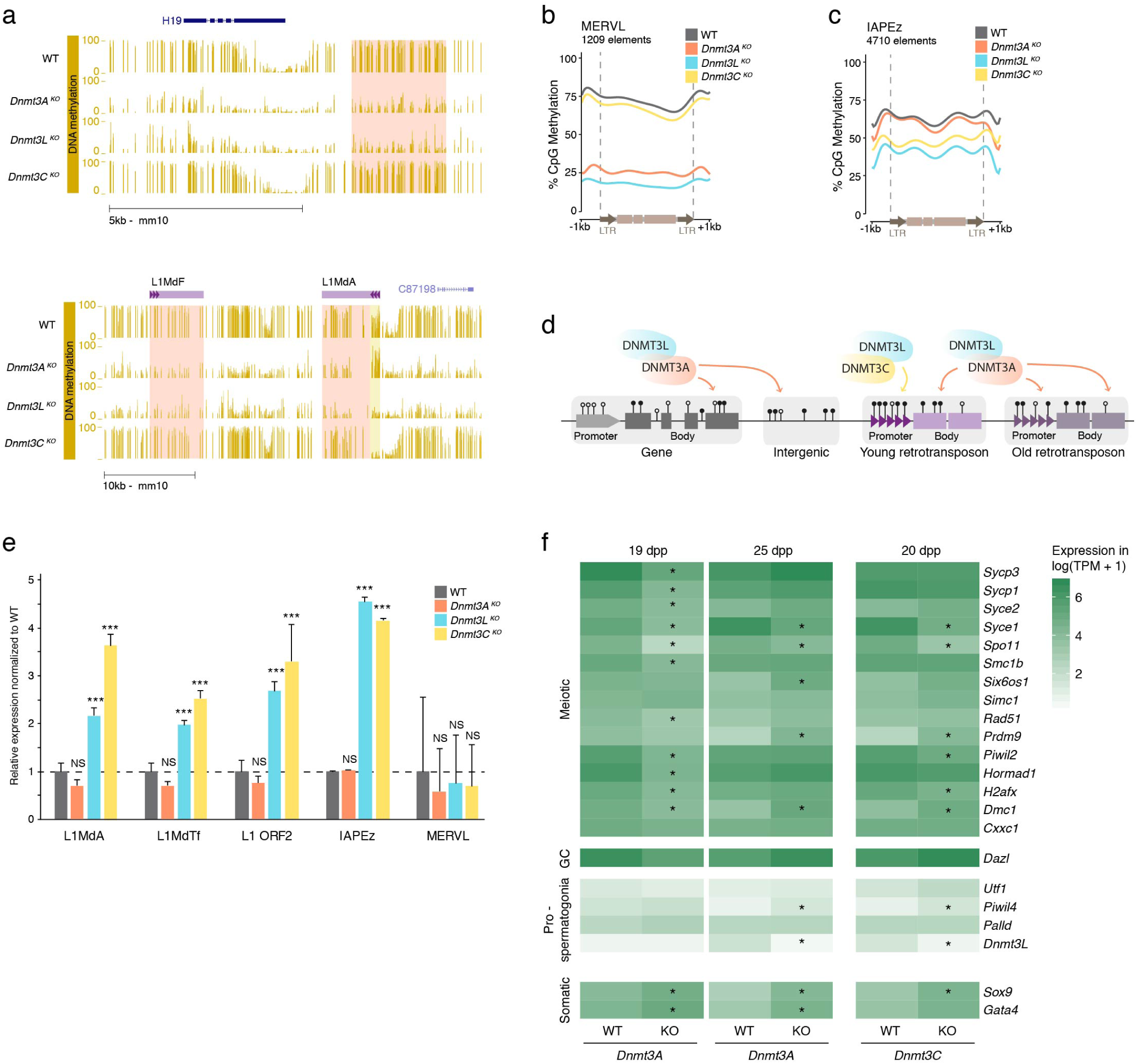
DNMT3A is not required for retrotransposon silencing. **a**, Genome browser representation showing DNA methylation levels assessed by WGBS at E18.5 in WT,*Dnmt3A^KO^, Dnmt3L^K0^* and *Dnmt3C^K0^* prospermatogonia. Top panel:the highlighted region (pink) coincides with the imprinting control region of the H19-lgf2 locus,with decreased methylation in *Dnmt3A^KO^* and *Dnmt3LKo.* Bottom panel: an evolutionary ancient L1MdF element is hypomethylated throughout its length in *Dnmt3A^KO^* (pink),while a recent L1MdA element shows hypomethylation over its promoter in *Dnmt3C^K0^* (yellow) and over its body in *Dnmt3A^KO^* (pink). **b and c**, Metaplots of DNA methylation levels over full-length MERVL (b) and IAPEz (c) comparing WT, *Dnmt3A^KO^, Dnmt3LKo,* and *Dnmt3CK ^0^.* IAP methylation levels are less decreased compared to other retrotransposon families,in agreement with a subset of IAPEz copies being resistant to DNA methylation reprogramming in PGCs^59^. **d,** Model of division of function for de novo DNA methylation in fetal male germ cells. DNMT3C selectively methylates the promoters of evolutionarily young retrotransposons. DNMT3A methylates the rest of the genome, including the body of young retrotransposons and the entire length of old retrotransposons. DNMT3L is a universal co-factor that stimulates both activities. **e**, Fold change expression (KO/WT) of different retrotransposon families in 19dpp whole testes, assessed by RT-qPCR normalized by beta-actin expression. Data are mean ± normalized SD (black bar) from at least three biological replicates, (student t-test over WT, *p<0.05, **p<0.005, ***p<0.0005). **f,** RNA-seq expression in log(TPM+1) (TPM, transcript counts per million) of genes specifically expressed during meiosis in prospermatogonia in 19dpp and 25dpp *Dnmt3A^KO^* and WT testes and in public datasets from 20dpp *Dnmt3C­ Ko* and WT 12

**Extended data Fig. 2.**
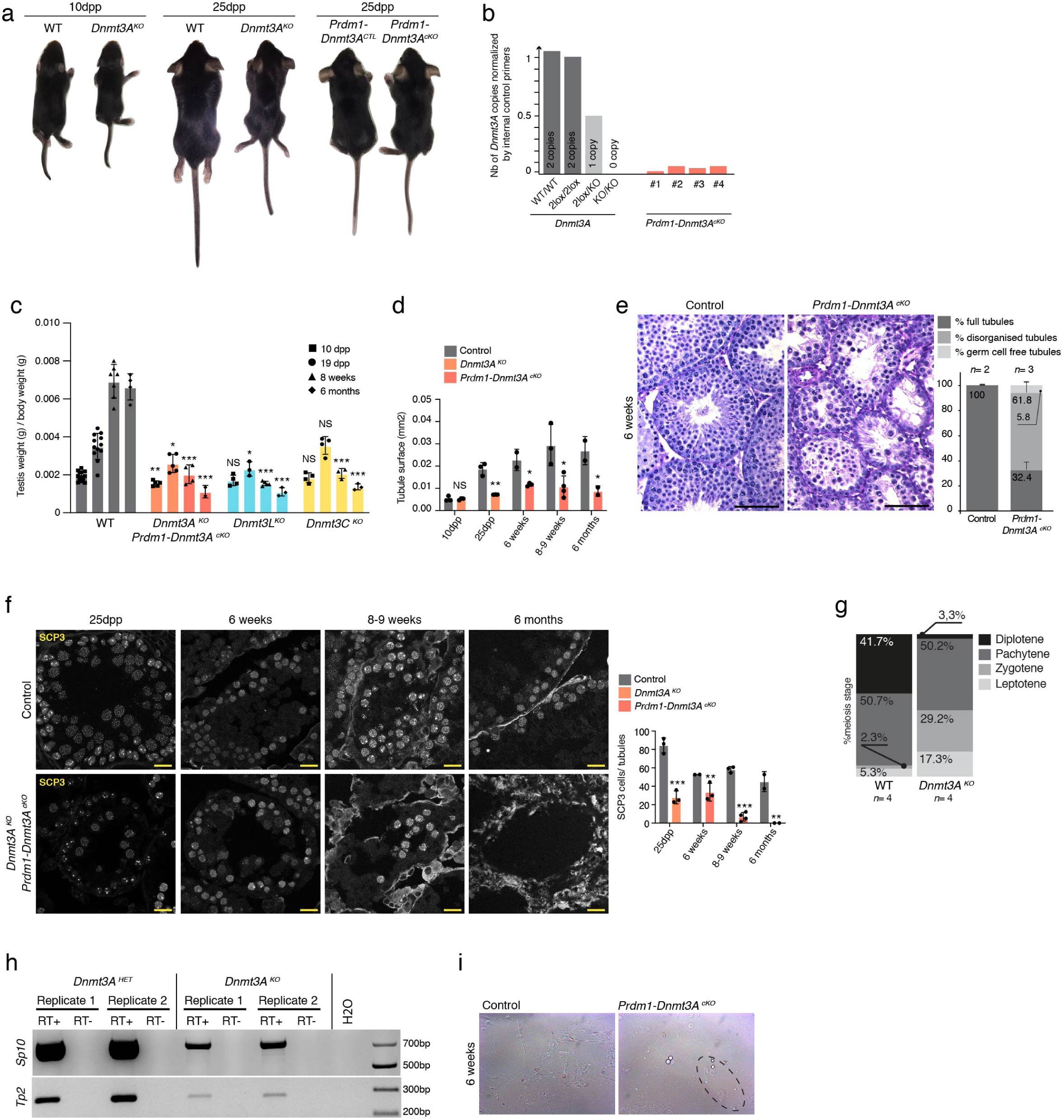
Dnmt3A mutants complete the first wave of spermatogenesis until spermatozoa with a reduced germ cell numbers. **a,** Representative photograph of a smaller constitutive *Dnmt3A^KO^* animal with its WT littermate at 1Oddp (left panel) and 25dpp (middle panel),and a germ cell-conditional *Prdm1-Dnmt3AcKo* animalwith its control littermate at 25dpp (right panel). All animals are males. **b,** Genomic qPCR of *Dnmt3A* copy number. *Dnmt3A^WT^, Dnmt3A2^10^ Dnmt3A^210^xlK ^0^, Dnmt3N°*correspond to liver DNA. Genomic DNA from *Prdm1-Dnmt3Ac^K0^* (# indicates individual replicates) was extracted from EpCAM-pos, [32M-neg FAGS-sorted germ cells (10dpp). Germ cell sorting purity assessed by TRA98 staining was the following:#2: 79%, #3: 96.8%, #4: 93.3%. c,Testicular weight normalized by body weight at 10dpp, 19dpp, 8 weeks and 6 months. Data are mean ± normalized SD (black bar) from biological replicates, and individual points represent biological replicates, (student t-test over WT, NS = non significant, *p<0.05, **p<0.005, ***p<0 .0005). **d,** Mean of tubule surface in mm2 in *Dnmt3A^KO^ Prdm1-Dnmt3N^K0^* and controls at 10dpp, 25dpp,6weeks, 8-9weeks and 6 months. Data are mean ± normalized SD (black bar) from biological replicates, and individual points represent biological replicates, (student t-test with p values as in c). e, (Left) Representative images of Periodic Acid Shift (PAS)-stained testis sections from germ cell-conditional*Prdm1-Dnmt3N^K0^* mutants and controls at 6 weeks (scale, 50 µm).(Right) Quantification of the percentage of different classes of tubules per genotype. Data are mean ± normalized SD (black bar) from biological replicates, n = number of animals. Control genotypes: *Dnmt3A^210^x1Ko; Prdm1-Cre^010^* and *Dnmt3A^KO/WT^; Prdm1-Crerg10_* **f,** (Right) Representative image of SCP3 (meiotic marker) staining on testis sections from *Dnmt3A^KO^, Prdm1-Dnmt3N^K0^* and controls at 25dpp, 6 weeks, 8-9weeks and 6 months (scale, 20 µm). (Left) Quantification of SCP3-positive cells normalized per tubule. Control genotypes at 6 weeks: *Dnmt3A^KO/WT^; Prdm1-CreTg/O* and *Dnmt3A21ox11<o; Prmd1-Cre^0/0^* - at 8-9 weeks: *Dnmt3N^10^x11<^0^; Prmd1-Cre^0/0^* (n= 2) and *Dnmt3N1oxNVT; Prdm1-Cre^0/0^* - at 6 months: *Dnmt3A^KO/WT^; Prmd1-Crerg/O* and *Dnmt3A^2lox/WT^; Prmd1-Cre^0/0^* Data are mean ± SD (black bar), individual points represent biological replicates (student t-test over WT *p<0.05, **p<0.005, ***p<0.0005). g, Percentage of the different stages of the prophase of first meiosis assessed by double-immunofluorescence detection of SCP3 and yH2AX on WT and *Dnmt3A^KO^* (25dpp) meiotic spreads (representative image on Fig. 2e). Data are mean for biological replicates, n = number of animals, at least 100 cells were counted per replicate. h, RT-PCR detection of spermatid-specific markers (Sp10 and Tp2) in *Dnmt3A^KO/WT^* and *Dnmt3AKD1^K0^* whole testes (25dpp). Two biological replicates for each genotype. i, Representative photograph of spermatozoa collected after epididymis squeezing in a conditional *Dnmt3N^K0^* mutant and control littermate *(Dnmt3A^KO/WT^; Prdm1-Crerg/O)* at 6 weeks

**Extended data Fig. 3.**
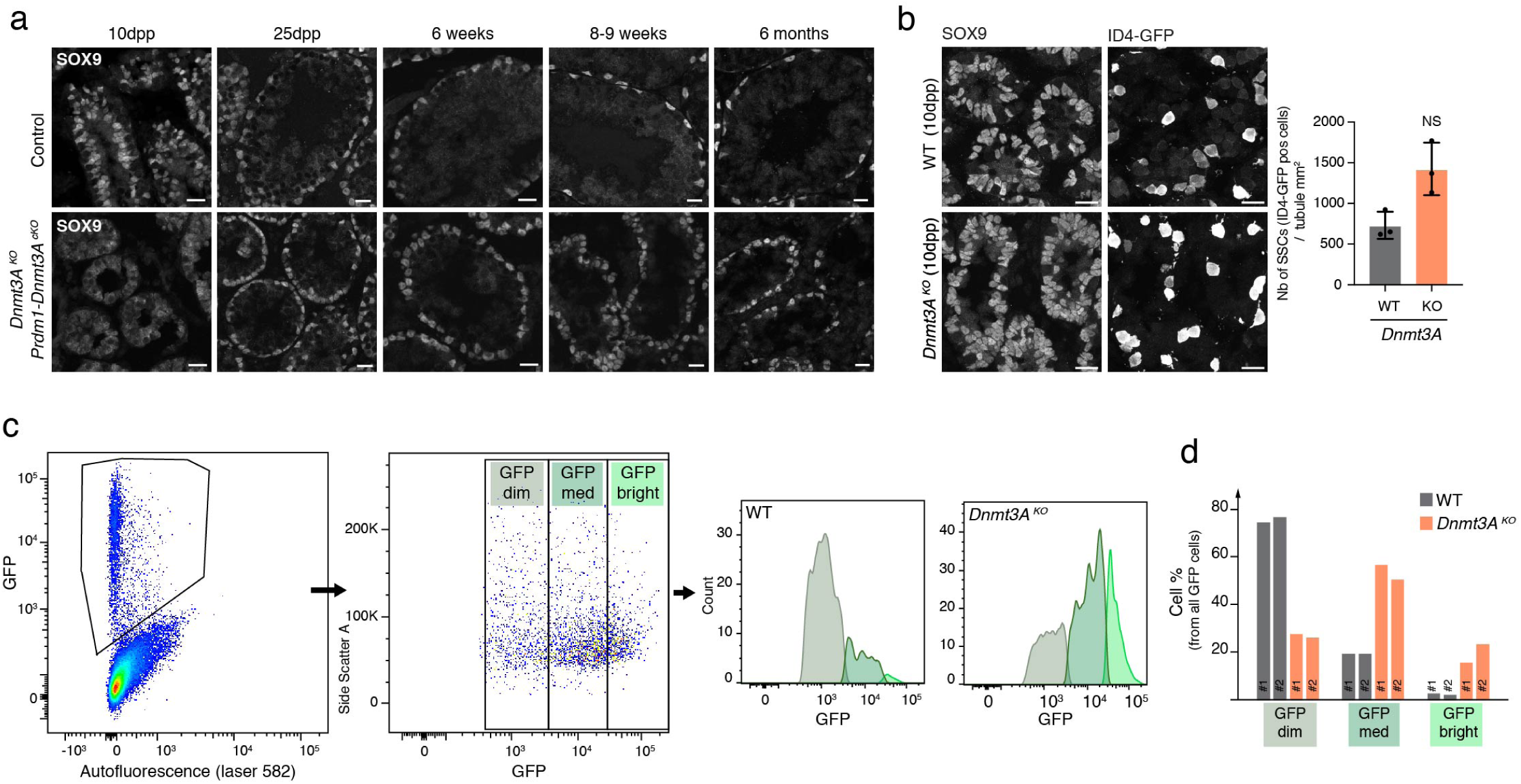
Accumulation of 104-eGFP positive germ cells in Dnmt3A^KO^ testes. **a**, Representative image of SOX9 (Sertolicell marker) staining on testis sections from *Dnmt3A^KO^, Prdm1-Dnmt3N^K0^* and controls at 10dpp, 25dpp, 6 weeks, 8-9weeks and 6 months. Scale, 20 µm. b, (Left) Representative microscopy images of double-immunofluorescence detection of SOX9 (Sertoli cell marker) and ID4-eGFP (SSC) at 1Odpp. Scale, 20 µm. (Right) Quantification of the number of ID4-eGFP positive cells per mm2 of tubule in *Dnmt3A^KO^* and WT. Data are mean ± SD (black bar) and individual points represent biological replicates (student t-test over WT, NS: non-significant). c, (Left) Representative FACS analysis plots of live ID4-eGFP-positive gated testicular cells from 1Odpp mice. ID4-eGFP cells were divided into three classes: GFP dim, GFP med and GFP bright. (Right) Representative count of cell numbers in each GFP subclass in *Dnmt3A^KO^* and WT d,Cell count in percentage for each GFP sub class in WT and *Dnmt3A^KO^* for two biological replicates. #1 et #2 represent individual replicates.

**Extended data Fig. 4.**
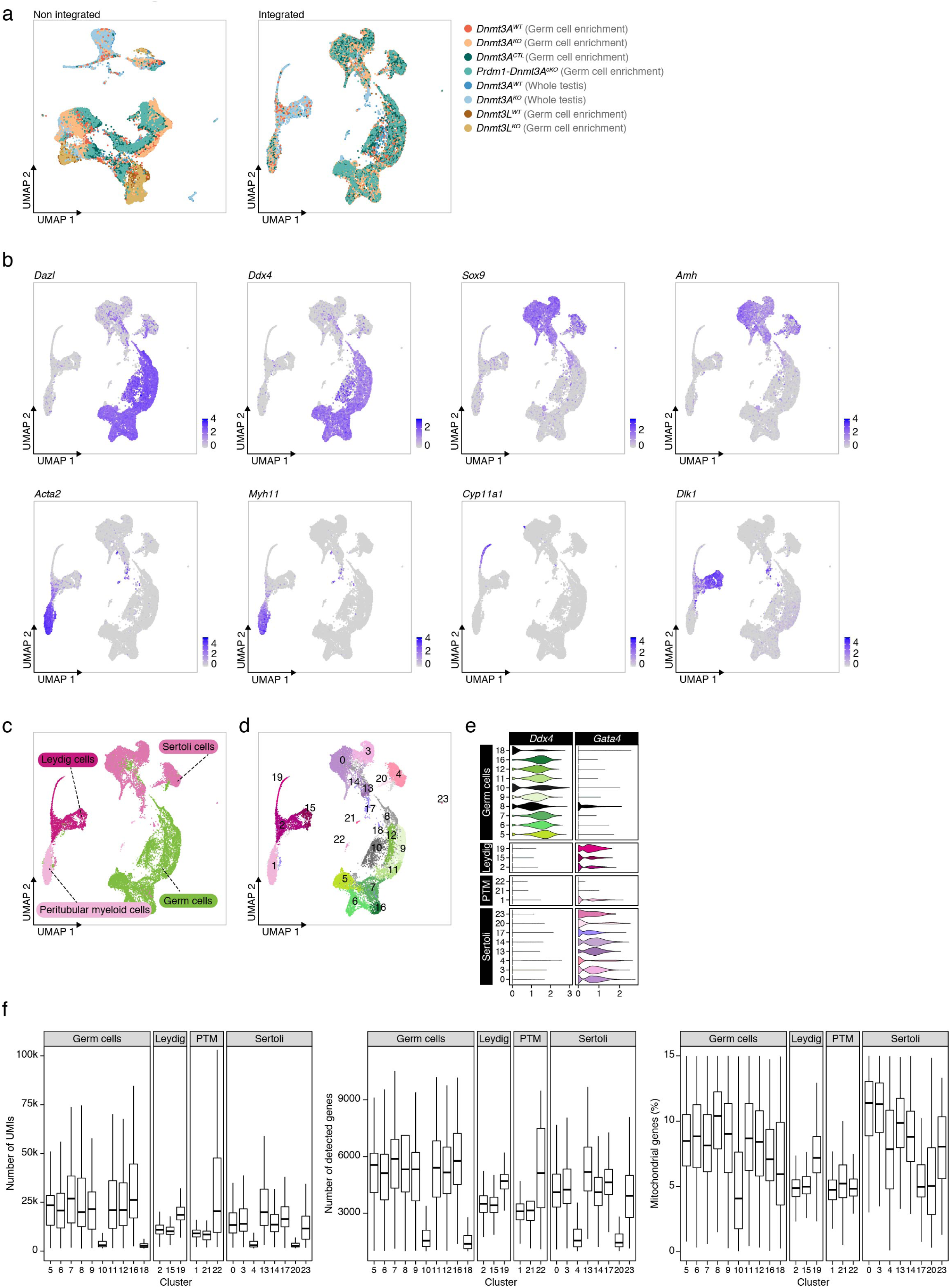
scRNA-seq quality controls and cluster annotation of germ and somatic cells. **a,** (Left) UMAP dimensionality reduction representation of scRNA-seq data from 1Odpp testicular cells before batch effect correction of eight samples, each sample is one animal of the genotype of interest (41,582 cells).Colors represent different conditions,see key.(Right) UMAP dimensionality reduction representation of the same sample after batch effect correction (see Methods, integrated data). **b,** UMAP representation of scRNA-seq data from 10dpp testicular cells *(n=* 8 animals from different genotypes) with a color gradient representing the expression of known markers of germ cells (Dazi and Ddx4), Sertoli cells (Sox9 and Amh), peritubular myeloid cells (PTM) (Myh11and Acta2) and Leydig cells (Dlk1 and Cyp11a1).c, UMAP representation of scRNA-seq data from 1Odpp testicular cells, colors represent cellular types based on marker expression of b. **d,** Unbiased cell clustering onto an UMAP representation of scRNA-seq data from 10dpp testicular cells demonstrated 24 clusters, each color represents a cluster annotated by a number (from 0 to 23). **e,** Violin plots show expression of key cell-type specific markers among cell clusters to distinguish those containing germ cells (Ddx4-positive) from testicular somatic cells (Gata4-positive). Cluster 8 presented expression of the two markers: this germ cellcluster was considered as somatically contaminated-probably due to cellular doublets-and was removed from further analysis.Cluster in black was removed from analysis because of poor quality control (see Extended data Fig.4f).**f,** Quality control of scRNA-seq integrated data split by cellular cluster, presenting the number of UMls {left panel), the number of detected genes (middle panel) and the percentage of mitochondrial reads (right panel).

**Extended data Fig. 5.**
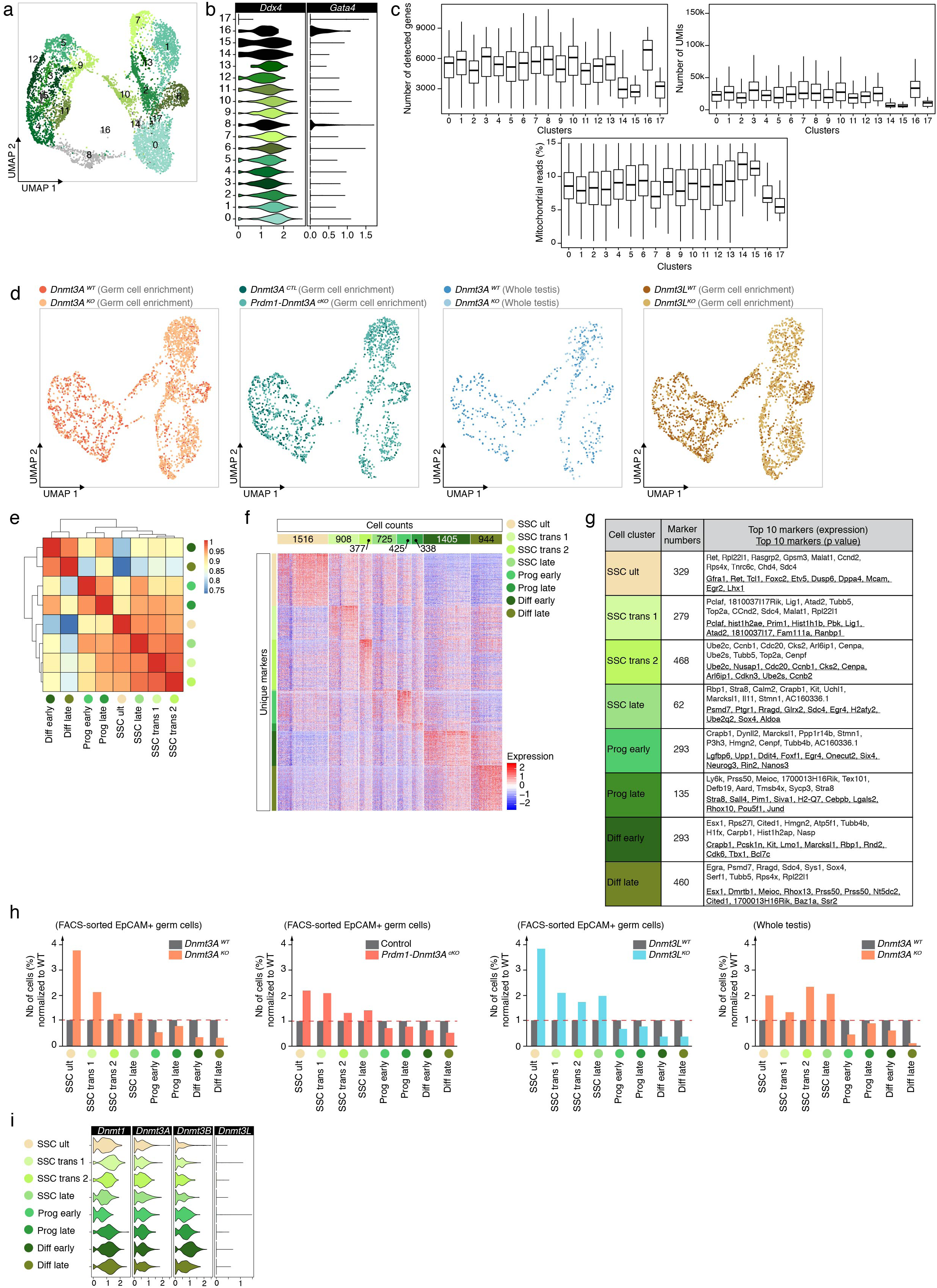
Supporting evidence to name final germ cell clusters. **a**, Unbiased cellclustering onto an UMAP representation of scRNA-seq data from 10dpp germ oells (n = 8 independent animals from different genotypes). Germ cells were selected from clusters 5, 6, 7, 9, 11, 12, 16 based on Dxd4 expression and quality controls. Identification of 18 clusters (from 0 to 17), colors represent the different clusters, cluster in grey (#8) was removed from final analysis (see Extended data Fig. 5b,c). **b,** Violin plots show expression of key cell-type specific markers among germ cell clusters to distinguish those containing germ oells (Ddx4-positive) from testicular somatic cells (Gata4-positive). The five clusters in black were removed from further analysis: clusters 14, 15 and 17 because of poor sequencing quality (Extended data Fig. 5c), and clusters 8 and 16 because of somatic contamination probably due to cellular doublets. **c**, Quality control of scRNA-seq integrated data on germ cell clusters. Cluster 14, 15 and 17 were removed from further analysis for poor sequencing quality. **d,** UMAP dimensionality reduction representation of scRNA-seq integrated data from 10dpp germ cells (6,638). Colors represent the different samples and conditions,red:*Dnmt3A^WT^* FACS-enriched germ cells, orange: *Dnmt3A^KO^* FACS-enriched germ cells,dark green: *Dnmt3A^CTL^ (Dnmt3A^2lox/KO^; Prdm1-Cre^0/0^)* FACS-enriched germ cells, light green: *Prdm1-Dnmt3N^K0^ (Dnmt3A^2lox/KO^; Prdm1-Cre^Tg/0^)* FACS-enriched germ cells, dark blue: *Dnmt3A^WT^* all testicular cells, light blue: *Dnmt3A^KO^* all testicular cells, brown:*Dnmt3L^WT^* FACS-enriched germ oells, yellow: *Dnmt3L^KO^* FACS-enriched germ cells. e, Heatmap of pairwise Spearman correlation matrix among the 8 germ cell clusters.**f,** Heatmap of single cell expression profile of markers defined in only one cluster for each germ cell cluster. Numbers on the upper row represent the number of cells in each germ cell cluster. **g,** Table of top 10 markers for each germ cell cluster. Marker numbers correspond to the total number of identified markers that give a signature for each cluster. **h,** Bar plot showing the percentage of cells normalized to *WT* per cell type for each mutant and condition.i, Violin plots show expression of Dnmt3 genes across germ cell populations.

**Extended data Fig. 6.**
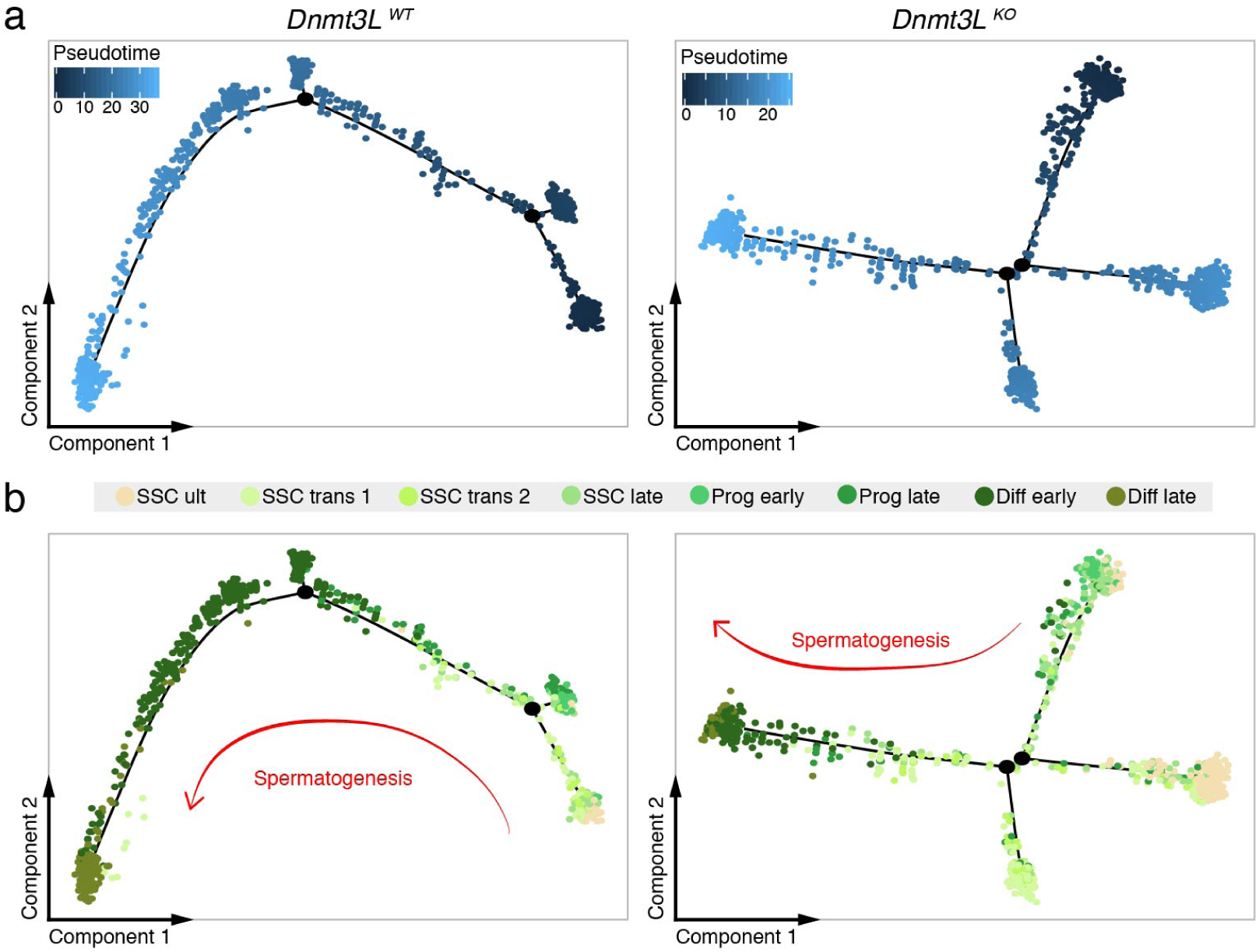
IDNMT3L is essential to SSC plasticity.a, b, Pseudotime trajectory of germ cells from 10dpp testes. (Left) *Dnmt3L^WT^* germ cells (FACS-sorted). (Right) *Dnmt3L^KO^* germ cells (FACS-sorted). In (a), cells were ordered from beginning (dark blue) to the end (light blue). In (b), cells were colored following germ cell cluster allocation, see key above·. Unfortuna­ tely, RNA velocity could not be performed on *Dnmt3L^KO^,* due to insufficient germ cells (1,031 WT and 1,182 *Dnmt3L^KO^*) and the use of anterior version of the scRNA-seq kit (10X Genomics V2).

**Extended data Fig. 7.**
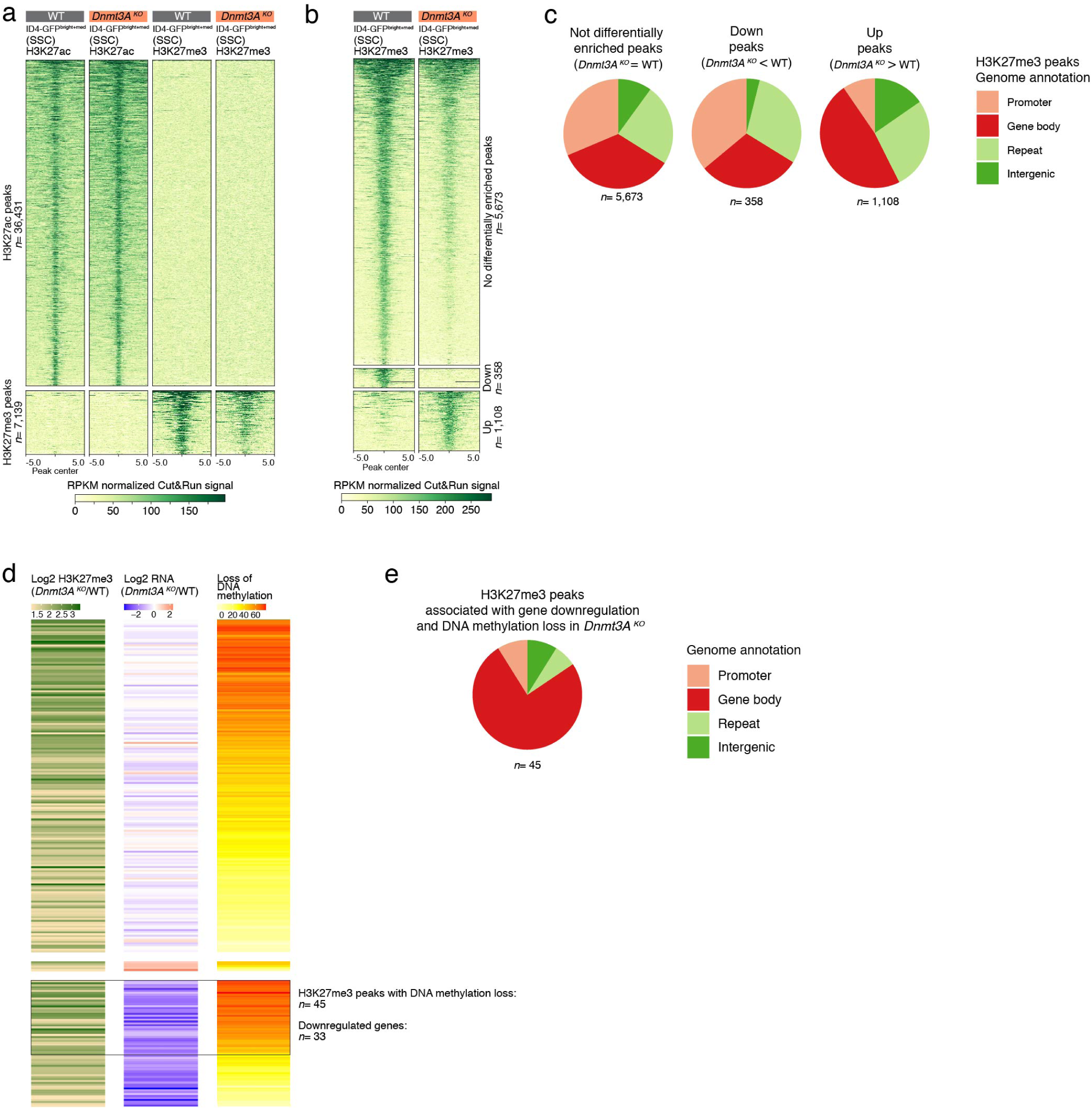
Relationship between H3K27me3 distribution, gene expression and DNA methylation in *Dnmt3A^KO^* SSCs (ID4-eGfp^bright+med^). **a,** Heat map showing levels of H3K27ac and H3K27me3 peak enrichment in RPKM normalized Cut&Run signal for ID4-eGFP^bright+med^ germ cells (SSCs) from 10dpp *Dnmt3A^KO^* and WT males. Enrichment is assessed +/- 5kb from the center of the peak. Number of biological replicates:*Dnmt3N°=* 3,WT= 4.There is no reciprocal gain or loss of one lark at the expense of the other *Dnmt3A^KO^* germ cells. b, Heatmap showing levels of H3K27me3 peak enrichment in RPKM normalized Cut&Run signal for ID4-eGFP^bright+med^ germ cells (SSCs) in 10dpp *Dnmt3N°* and WT males. Peaks are divided into three categories: not differentially enriched *(Dnmt3A^KO^* = WT), down-enriched *(Dnmt3A^KO^* < WT), up-enriched *(Dnmt3N°*> WT) (FDR<5% and FC>1). c, Genomic annotation of each category of H3K27me3 peaks. d, Heatmap focusing on i) *Dnmt3A^KO^-gained* H3K27me3 peaks (log2 FC) compared to WT (left), ii) differential expression (log2 FC) of genes associated with H3K27me3 peaks (TSS +/- 5kb from the peak) between *Dnmt3N°* and WT-from ID4-eGfp^bright+med^ RNA-seq-(center), and iii) percentage of DNA methylation loss in *Dnmt3A^KO^* versus WT on H3K27me3 peak genomic location-from E18.5 WGBS-(right). Rows are ordered according to gene expression changes: top, not differentially expressed; middle, upregulated; bottom, downregulated. e, Genomic annotation of *Dnmt3A^KO^-gained* H3K27me3 peaks, associated with downregulated genes and >30% DNA methylation loss on H3K27me3 peak location.

**Extended data Fig. 8.**
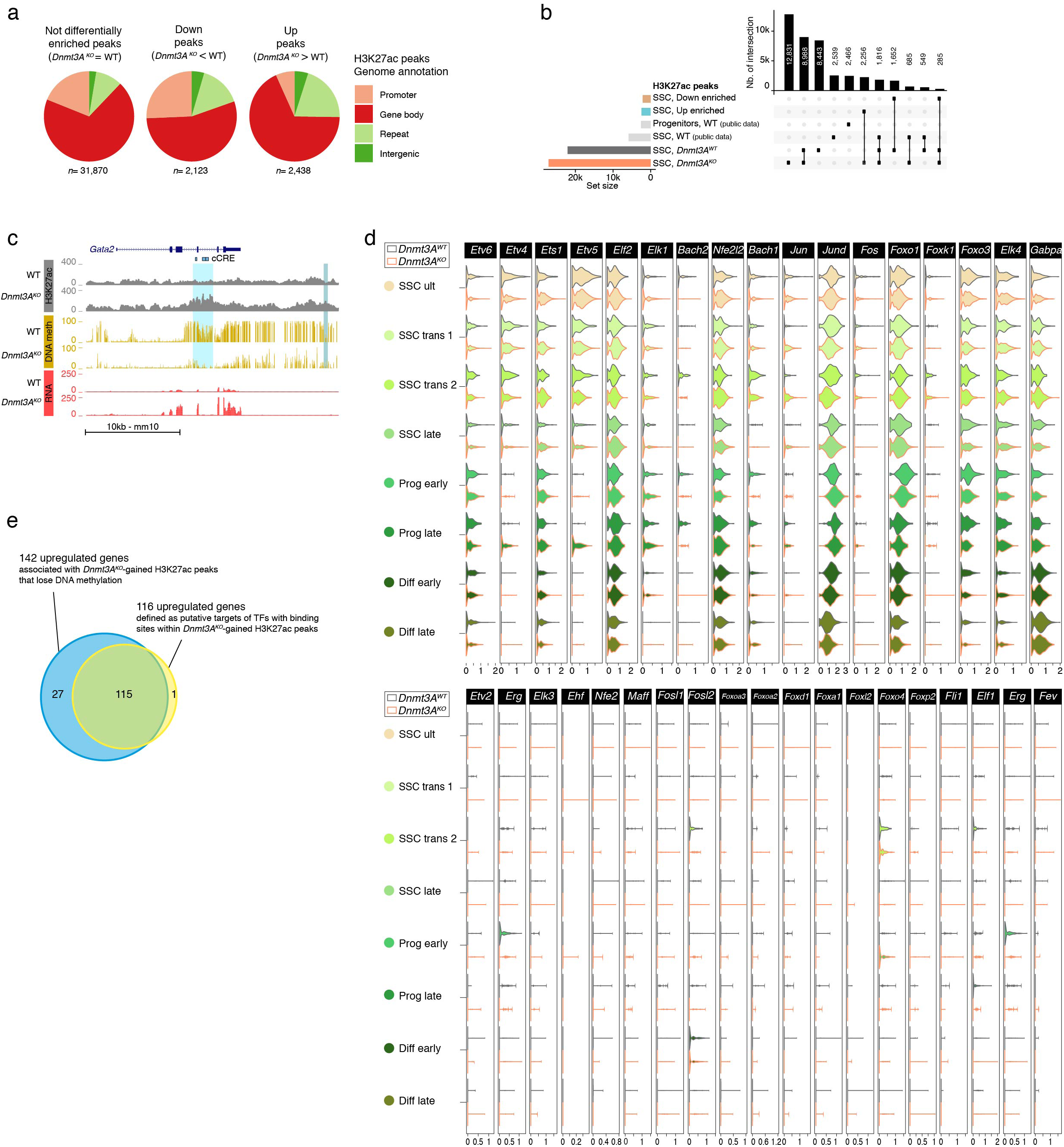
Relationship between H3K27ac distribution, gene expression and DNA methylation in Dnmt3A^KO^ SSCs (ID4-eGFP^bright+med^). **a,** Genomic annotation of each category of H3K27ac peaks. **b,** Quantification of intersection between H3K27ac peaks from *Dnmt3A^K0^, Dnmt3A^WT^.* WT SSC (ID4-eGFP^bright^) and progenitors (ID4-eGFP^dim^) (from published data^30^), down-enriched peaks and up-enriched peaks. Intersections lower than 200 were removed from the figure.c,Genome browser representation of the Gata2 locus, showing H3K27ac enrichment (grey), DNA methylation (yellow) and RNA expression (red) for *Dnmt3N°*and WT. Regions showing *Dnmt3A^KO^* de nova H3K27ac peaks associated with >30% DNA methylation loss are shaded in light blue; regions showing de nova H3K27ac peaks at regions not controlled by DNA methylation are shaded in grey-blue. **d,** Violin plots showing expression levels of TF factors that have the best match with the motifs present in Dnmt3A^KO^-specific H3K27ac peaks. Expression is represented across SSCs and spermatogonia subtypes discriminated by scRNA-seq for *Dnmt3A^KO^* and WT. Upper panel: 17 TFs with SSC/spermatogonia expression, lower panel: 18 TFs not expressed in SSC/spermatogonia. e, Venn diagram representation of the overlap between 142 upregulation genes associated with *Dnmt3A^KO^-gained* H3K27ac peaks that lose DNA methylation (see Fig. Se), and 116 upregulated genes defined as putative targets of TFs with binding motif sites within *Dnmt3N°-gained* H3K27ac peaks (see Fig.Si).

